# The early endosomal protein Rab21 is critical for enterocyte functions and intestinal homeostasis

**DOI:** 10.1101/2020.11.18.378984

**Authors:** Sonya Nassari, Dominique Lévesque, François-Michel Boisvert, Steve Jean

## Abstract

Membrane trafficking is defined as the vesicular transport of molecules into, out of, and throughout the cell. In intestinal enterocytes, defects in endocytic/recycling pathways result in impaired function and are linked to genetic diseases. However, how these trafficking pathways regulate intestinal tissue homeostasis is poorly understood. Using the *Drosophila* intestine as an *in vivo* model system, we investigated enterocyte-specific functions for the early endosomal trafficking machinery in gut homeostasis. We focused on the small GTPase Rab21, which regulates specific steps in early endosomal trafficking. Rab21-depleted guts showed severe abnormalities in intestinal morphology, with deregulated homeostasis associated with a gain in mitotic cells and increased cell death. Increases in both apoptosis and yorkie signaling were responsible for compensatory proliferation and tissue inflammation. Using a RNA interference screen, we identified specific regulators of autophagy and membrane trafficking that phenocopied Rab21 loss. We further showed that Rab21-induced hyperplasia was rescued by inhibition of epidermal growth factor receptor signaling, and identified improperly trafficked cargoes in Rab21-depleted enterocytes. Our data shed light on an important role for early endosomal trafficking, and particularly Rab21, in enterocyte-mediated intestinal homeostasis.

## INTRODUCTION

Membrane trafficking is characterized by the vesicular transport of proteins and macromolecules throughout the various cellular compartments, as well as in and out of cells. This process requires exchanges between compartments that are essential to maintain cellular functions and help cells adapt to the ever-changing conditions in the extra- and intra-cellular environments. The intricate link between membrane trafficking and cellular signaling means that modulating trafficking has strong consequences on cellular homeostasis [1]. Mutations in membrane trafficking genes are associated with a large array of human diseases [2]. Importantly, some of these mutations affect several tissues, while others are restricted to one organ system, and can be used to shed light on the cell typespecific functions of vesicular trafficking genes.

The digestive tract performs several essential functions, including food digestion, nutrient/water absorption, and protection against external pathogens. These tasks are achieved by enterocytes, a differentiated cell population constituting approximately 80% of the intestinal epithelium. Given their absorptive function, enterocytes display high membrane trafficking flux, which must be tightly orchestrated to ensure appropriate organ activity [3]. Defects in trafficking processes affect enterocyte properties (e.g., their shape, absorptive function, and resistance to stress) and lead to intestinal diseases [3,4]. Intestinal pathologies, such as microvillus inclusion and chylomicron retention disease, are associated with mutations in the membrane trafficking related-genes myosin 5B (*MYO5B*) and secretion associated Ras related GTPase 1B (*SAR1B),* respectively [2,5,6]. MYO5B is required for proper recycling to the apical membrane [5] while SAR1B is necessary for chylomicron secretion [6].

The small Rab GTPases constitute an important class of membrane trafficking regulators [7]. Rabs localize to specific cellular compartments, where they recruit various effectors to mediate their functions. In enterocytes, a few studies have reported the involvement of Rab8 and Rab11, both components of the endosomal recycling machinery, in the regulation of apical brush border formation [3,4,8]. In *Rab8-knockout* mice, apical peptidases and transporters accumulate in lysosomes. This leads to decreased absorptive functions, which are associated with microvillus inclusion and defects in microvillar structure [4]. Similarly, loss of *Rab11* in mouse intestinal epithelial cells results in defects in enterocyte polarity and abnormal microvilli organization [8]. Notably, the enterocyte polarity defect upon Rab11 knockdown is conserved in *Drosophila* [9]. In the *Caenorhabditis elegans* intestine, Rab11 also displays an apical distribution, which is required to establish proper polarity in intestinal epithelial cells [10]. Other studies have demonstrated roles for Rab11 in toll-like receptor regulation and microbe tolerance in mouse and *Drosophila* intestines [11]. Finally, recent work performed in mice and flies highlighted a conserved role for Rab11 in intestinal tumor progression, through the regulation of the Hippo pathway [9,12]. Although roles for the endosomal recycling machinery in enterocytes have been well documented, little is known about the functions of other membrane trafficking routes in these cells, and more generally, in intestinal tissue homeostasis.

Early endosomes act as sorting centers, in which endocytosed cargos must be correctly directed toward either recycling routes or lysosomal degradation [13]. Therefore, early endosomes are important for the proper regulation of internalized signals [1]. Rab5 is often associated with early endosomes, where it acts in endosomal homotypic fusion and in endocytosis [14]. Rab5 is also necessary for phosphatidylinositol-3-phosphate (PtdIns(3)P) synthesis in early endosomes [15]. In flies, Rab5 is required for the proper localization of apical proteins in polarized epithelia [16,17], while in mice, it is required for lysosome homeostasis in the liver [18]. Several other Rabs are also present in early endosomes [19], including Rab21, which is involved in endocytosis and early endosomal trafficking of integrins, epidermal growth factor receptor (EGFR), and certain clathrin-independent cargos [20–23]. Rab21 was first identified in human intestinal Caco-2 cells [24] and is expressed mainly in enterocytes in both human and mouse intestines [24,25]. Interestingly, recent studies demonstrated that Rab21 expression is decreased in enterocytes in solute carrier family 15 (oligopeptide transporter), member 1 (*Slc15a1)-knockout* mice [25]. *Slc15a1* is a susceptibility variant gene for inflammatory bowel disease (IBD) [26,27].

These data led us to investigate the specific functions of Rab21 and other trafficking regulators in enterocytes *in vivo* using the *Drosophila* intestine as a model system. *Drosophila* enables the use of powerful genetic tools, and is an excellent model to investigate intestinal biology, with intestinal signaling pathways and functions highly similar to those of humans [28]. Like the mammalian intestine, the *Drosophila* gut is composed of intestinal stem cells, which differentiate into progenitor cells that yield differentiated enteroendocrine cells and enterocytes [28–30]. The coordination of several signaling pathways orchestrates the balance between intestinal stem cell proliferation, progenitor differentiation, and differentiated cell turnover. Under normal conditions, Wnt signaling is required for intestinal stem cell self-renewal, similar to in mammals [31,32], while high levels of Notch signaling in progenitor cells promotes enterocyte differentiation [29,30]. Upon stress, the Janus kinase-signal transducer and activator of transcription (JAK-STAT), EGFR, c-Jun NH(2)-terminal kinase (JNK), and Hippo signaling pathways activate intestinal stem cell proliferation and progenitor differentiation in response to various injuries [33]. Furthermore, upon intestinal epithelial stress, enterocytes serve as stress sensors to initiate tissue regeneration, and rapidly compensate for cell loss by promoting the non-cell autonomous activation of intestinal stem cells. The JNK and Hippo/yorkie (yki) pathways are involved in this process in enterocytes, where they trigger expression of the proinflammatory cytokine unpaired 3 (upd3) [34–39]. Secretion of upd3 promotes intestinal stem cell proliferation by directly activating JAK-STAT and indirectly the EGFR pathway [34,39,40].

Here, we demonstrate that functional early endosomes are required to maintain proper tissue homeostasis in fly intestines. Loss of Rab21 in *Drosophila* enterocytes results in yki activation and apoptosis, which both induce secretion of the pro-inflammatory cytokine upd3, leading to compensatory proliferation in a non-cell autonomous manner. An enterocyte focused screen of membrane trafficking genes reveals that blocking autophagy, endosomal trafficking, and EGFR-mitogen-activated protein kinase (MAPK) signaling also induces tissue inflammation and compensatory proliferation. Using epistasis experiments, we show that Rab21 independently regulates enterocyte EGFR signaling and autophagy. Additionally, we highlight a previously unappreciated contribution of Rab21 in PtdIns(3)P and PtdIns(3,5)P2 regulation. Finally, using a tandem mass tag (TMT)-based quantitative proteomic approach we shed light on deregulation of SLC transporters, intestinal proteases and cytochrome P450 proteins in enterocytes depleted for Rab21.

## RESULTS

### Enterocyte Rab21 is required to maintain intestinal epithelium morphology

Fluorescence-activated cell-sorted and single-cell RNA-sequencing databases indicate that the early endosome components Rab5 and Rab21 and the recycling endosome component Rab11 are expressed in various cell types throughout the gut. Rab21 is normally expressed at lower levels than Rab5; however, it is upregulated upon infection [41–43]. As the role of Rab11 in gut homeostasis is well-described [3], we decided to focus our initial analysis on Rab21, which plays roles in specific early endosomal trafficking events [20–22,44,45] and is modulated by stress in both flies and in mouse models of IBD [25,41]. We first determined the precise Rab21 expression pattern in the gut using a GAL4 driver line with GAL4 controlled by the *Rab21* promoter [46] and a UASt-green fluorescent protein (GFP):Rab21 responder line. In accordance with transcriptomic data [41–43], we observed that Rab21 was present throughout the digestive tract (Figure 1A-B), with diverse expression levels between the three main regions of the midgut (Figure 1A-B). Interestingly, Rab21 displayed stronger expression in the distal part of the posterior midgut (R5 region) [41] and in the copper cell region (Figure 1B). In the R5 region, Rab21 expression was enriched in large polyploid cells with low levels of armadillo (arm) staining, characteristic of enterocytes (Figure 1C-D, arrowheads), with no restricted basal or apical localization (Suppl. Fig. 1A). Using *in situ* hybridization, we confirmed that Rab21 was expressed in enterocytes in the posterior midgut, which were visualized with 4’,6-diamidino-2-phenylindole (DAPI) to highlight their polyploid nuclei (Suppl. Fig. 1B). We also detected Rab21 in cells displaying high arm staining, which were likely intestinal stem cells (Figure 1C-D, arrows). Immunofluorescence against Delta (Dl) confirmed that Rab21 was present in intestinal stem cells (Dl^+^; Suppl. Fig. 1C). It was also detected in prospero (pros)^+^ cells (Suppl. Fig. 1C), revealing that enteroendocrine cells also express Rab21. As previously observed in human and mouse intestines, these data show that Rab21 is expressed throughout the fly gut, with high expression in enterocytes, suggesting potential important functions in these cells.

**Figure 1.**
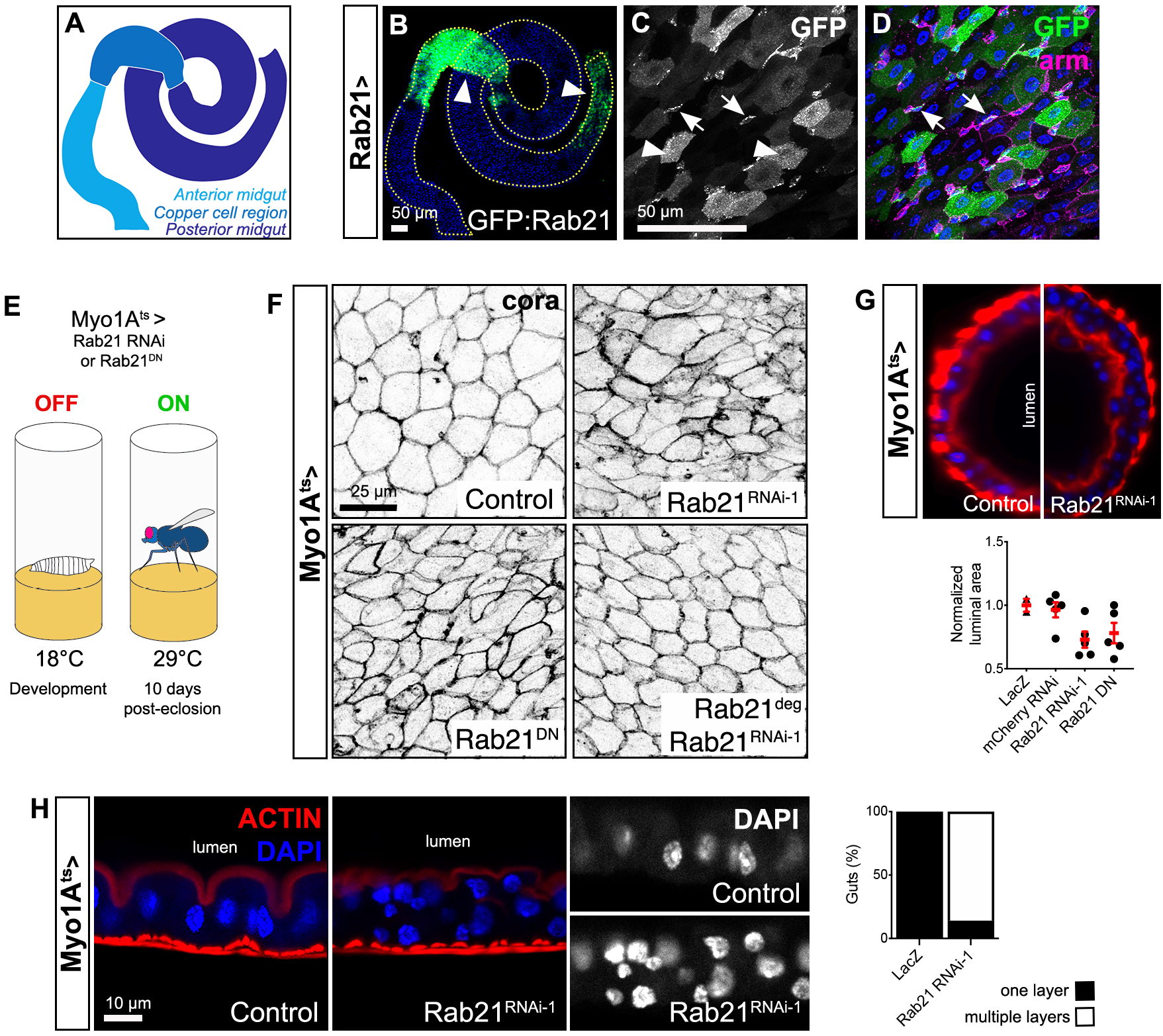
Rab21 expression in enterocytes is required to maintain correct intestinal epithelium morphology. (A-D) Schematics of *Drosophila* midgut compartmentalization and images of a *Drosophila* midgut expressing GFP:Rab21wt under the control of the *Rab21* promoter, using the Gal4-UAS expression system. (B) Blue is for DAPI staining and arrowheads indicate the copper cell region with the distal posterior midgut and (C-D) enriched Rab21 in enterocytes. (C-D) Cell junctions were immunostained for armadillo (arm). (C-D) Arrows show Rab21 enrichment in other arm^+^ cell types. Images are representative of *n*=15 guts from *N=3* independent experiments. (E) Experimental design. Using the temporally inducible Gal80^ts^ expression system, we independently induced one of two Rab21 RNAi hairpins or Rab21^DN^ expression in enterocytes. Flies were housed at 18°C during development and at 29°C for 10 days after eclosion prior to analysis. (F) Cell junctions were immunostained for coracle (cora) in control, Rab21-depleted enterocytes, and Rab21-depleted enterocytes overexpressing Rab21^deg^. *N*=3 independent experiments. (G) Transverse views of guts from control and Rab21-depleted flies stained with DAPI (blue) and phalloidin (red) (to detect actin). The normalized luminal area in *N*=2 independent experiments is shown. (H) Epithelial organization of guts from control and Rab21-depleted flies stained with DAPI and phalloidin. The percentages of guts with single- and multilayered epithelia are shown (*n*=14 guts from *N*=3 independent experiments).

To characterize Rab21 functions in enterocytes, we performed loss-of-function experiments using a cell-specific inducible system [47]. Two independent and previously validated [21] Rab21 RNA interference (RNAi) hairpins and a dominant negative (DN) Rab21 construct (Rab21^DN^) [21] were specifically expressed in the enterocytes of adult *Myo1A-Gal4, UAS GFP; tub-Gal80^ts^* (hereafter referred to as *Myo1A^ts^*) transgenic flies. Flies were housed at 18°C during embryonic development and at 29°C for 10 days after hatching, then analyzed (Figure 1E). Loss of Rab21 function was associated with a significant increase in cell density (Suppl. Fig. 2A), indicating defects in tissue morphology. Consistent with this notion, compared to controls (Figure 1F; Suppl. Fig. 2B), the organization of the cell junction markers coracle (cora; Figure 1F) and arm (Suppl. Fig. 2B) was disrupted with decreased Rab21 expression. Contrary to Rab11 defects [48], we did not observe noticeable changes in the microvillar structure or polarity of Rab21-depleted enterocytes (Suppl. Fig. 2C), and Smurf assays revealed no defects in tissue permeability in Rab21-depleted guts (Suppl. Fig. 2D). Furthermore, Rab21 knockdown in enterocytes led to a multilayered epithelium (Figure 1H; Video 2), which was never observed in control flies (Figure 1H, Video 1). Consistent with this observation, the intestinal lumen area tended to be reduced in flies depleted of Rab21 (Figure 1G). Importantly, overexpression of Rab21^DN^ in enterocytes had similar effects (Figure 1F-G; Suppl. Fig. 2A), supporting the specificity of the observed phenotypes. Survival experiments showed that Rab21 loss-of-function through either RNAi or Rab21^DN^ expression negatively affected lifespan (Suppl. Fig. 2E). Importantly, gut morphology defects observed with Rab21 RNAi were suppressed by overexpression of a RNAi-insensitive Rab21 construct referred to as Rab21 degenerated (Rab21^deg^; Figure 1F), confirming the specificity of the Rab21 RNAi reagents. Finally, we assessed whether Rab21 gain-of-function would inversely affect tissue morphology. Overexpression of a constitutively active form of Rab21 (Rab21^CA^) resulted in no changes in cell density (Suppl. Fig. 3A) and the cell junction marker arm was decreased (Suppl. Fig. 3B). This suggests that Rab21 is necessary for proper gut morphology, but not sufficient to drive strong phenotypes. Altogether, these results highlight the contribution of enterocyte Rab21 in the maintenance of gut morphology, as well as its requirement during aging.

### Enterocyte Rab21 maintains intestinal cellular homeostasis

Enterocytes are non-mitotic, terminally differentiated cells arising from progenitors produced by intestinal stem cells (Figure 2A). Surprisingly, while never observed in control guts (Figure 2B), Rab21 depletion from enterocytes resulted in supernumerary cells underneath the Myo1A^+^ cells (Figure 2B). These cells did not express Myo1A, suggesting that they were not enterocytes. Furthermore, the percentage of Myo1A^+^ cells was decreased in guts depleted of Rab21 compared to control guts (Figure 2C). To identify the accumulating cells, we assessed the impact of Rab21 depletion in enterocytes on the other main midgut cell types. Upon Rab21 knockdown, we noticed increases in both the enteroendocrine cells and stem cell populations, using immunofluorescence against pros and Dl, respectively (Figure 2D, E). The increase in the proportion of intestinal stem cells was further confirmed using a Dl:GFP reporter line (Figure 2E). From these data, we conclude that enterocyte Rab21 is necessary for the maintenance of intestinal homeostasis.

**Figure 2.**
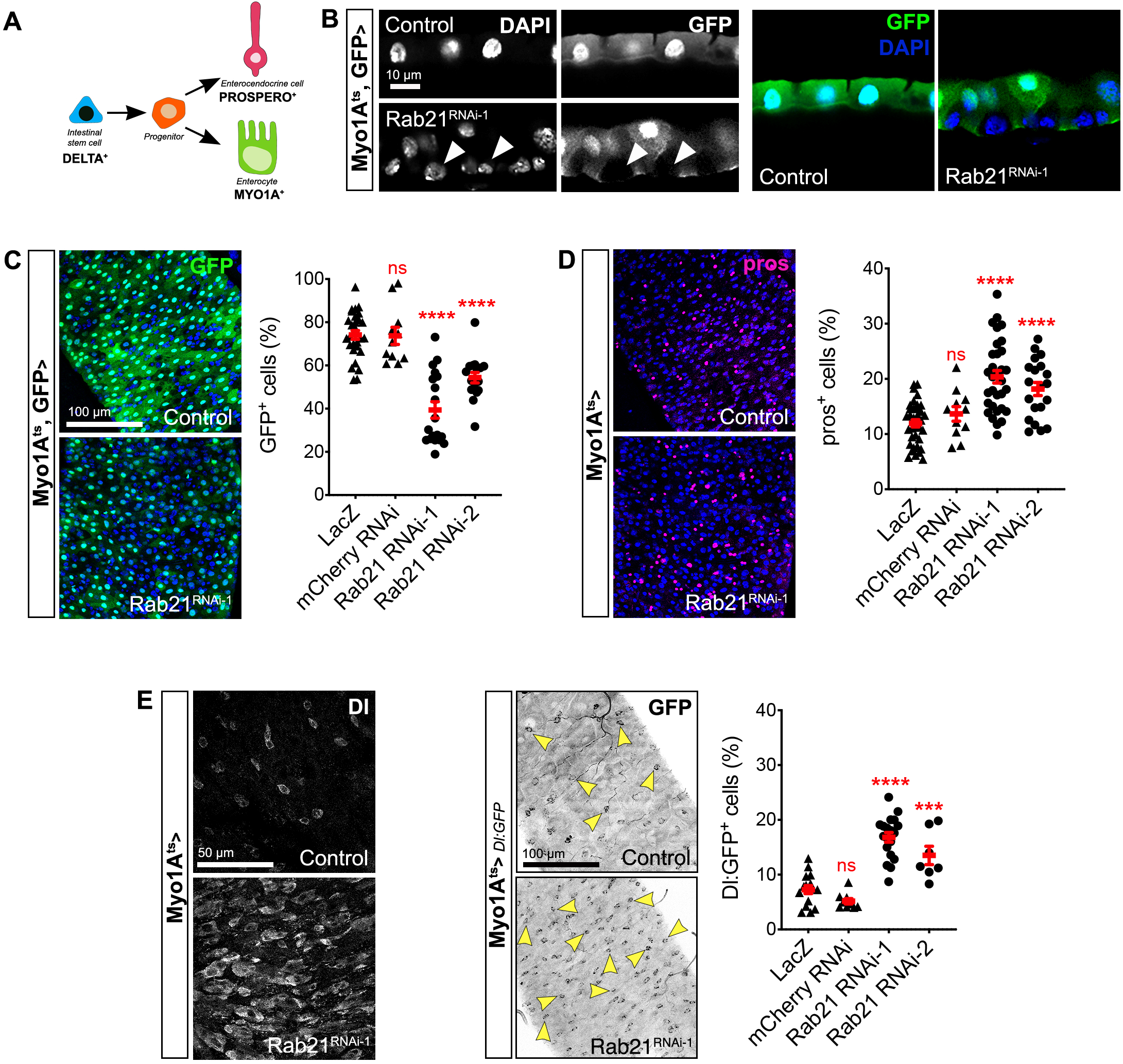
Enterocyte Rab21 is required to maintain intestinal homeostasis. (A) Schematic of cell lineages, markers, and drivers in the *Drosophila* gut. (B) GFP was expressed under the control of the enterocyte-specific *Myo1A* promoter to detect enterocytes in control and Rab21-depleted guts. Arrowheads indicate accumulating Myo1A^-^ cells. (C) Quantification of the percentage of Myo1A^+^ cells in control and Rab21-depleted guts in *N=3* independent experiments. (D) Rab21-depleted and control enteroendocrine cells were immunostained for prospero (pros). Cells positive for pros were quantified in *N*=3 independent experiments. (E) Intestinal stem cells were detected using Dl immunostaining or a Dl:GFP reporter. Arrowheads indicate Dl:GFP^+^ cells. Dl:GFP^+^ cells were quantified in *N*=3 independent experiments. *\*\*\*P* < 0.001; *\*\*\*\*P* < 0.0001 by unpaired *t*-test (C, D, E) or Mann-Whitney U test (E); ns, non-significant. All error bars are the SEM.

### Rab21 is necessary for proper cell turnover

Next, we investigated the signaling mechanisms impacting the various gut cell populations upon Rab21 depletion from enterocytes. In various contexts, increased cell density in the gut is compensated by increased apoptosis [37]. Hence, we first assessed cell death by immunostaining for cleaved caspase 3. In guts with Rab21-depleted enterocytes, the intensity of cleaved caspase 3 staining was significantly increased compared to controls (Figure 3A). The increased cell death in Rab21-depleted intestines was further corroborated using SYTOX staining (Figure 3B). Consistent with these results, the transcript level of head involution defective (hid), which promotes caspase activity, was upregulated upon Rab21 depletion (Suppl. Fig. 3C).

**Figure 3.**
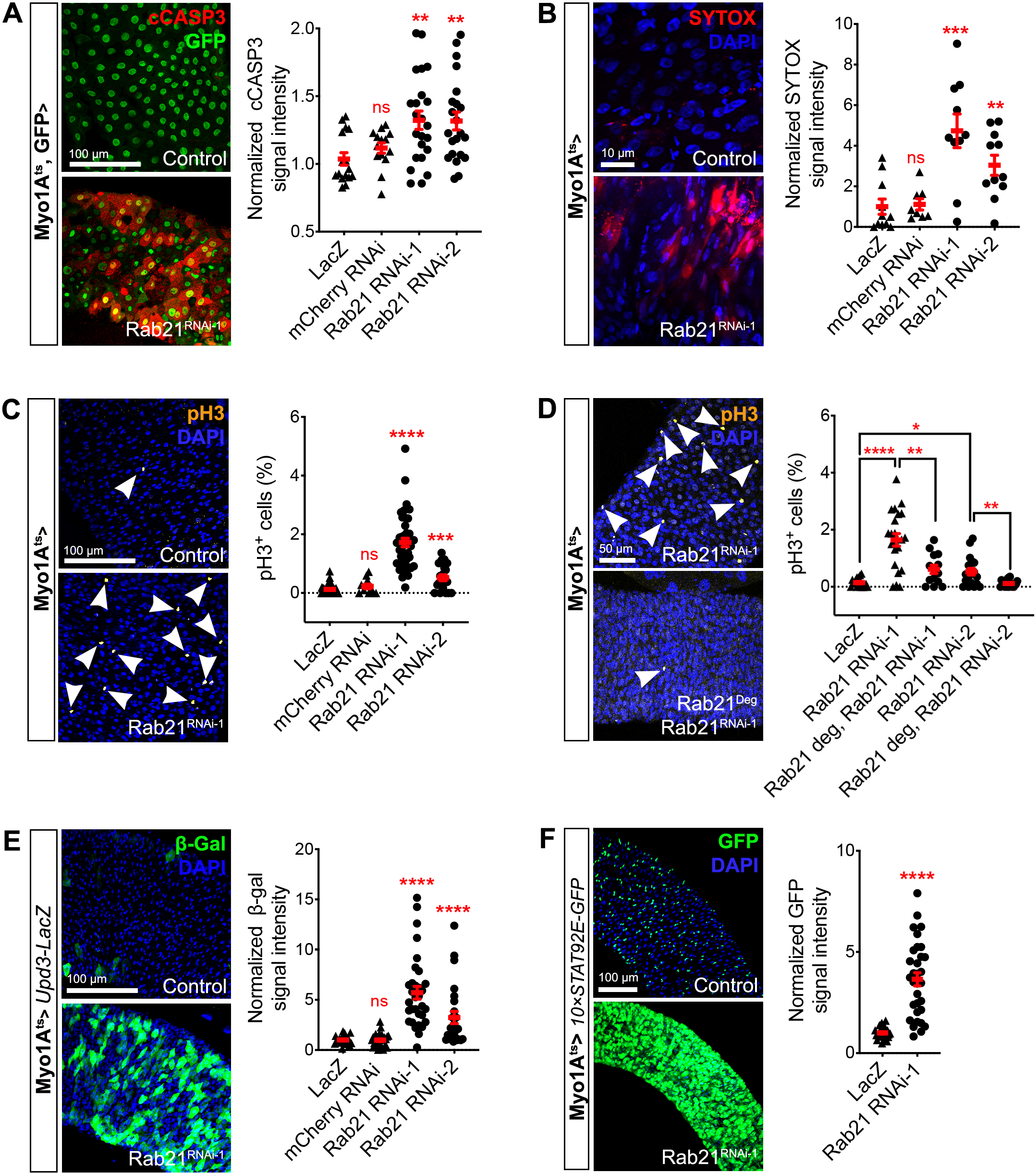
Rab21 is necessary for proper cell turnover. (A) Cleaved caspase 3 (cCASP3) immunostaining and (B) SYTOX staining were used to detect apoptotic cells. Quantifications of cCASP3 (A) and SYTOX (B) staining intensities are based on *N*=3 independent experiments. (C-D) Immunostaining for pH3 was used to detect mitotic cells. (C-D) Quantification of mitotic cell percentages was based on *N*=3 independent experiments. (E) *Drosophila* guts expressing a upd3-LacZ reporter were immunostained for β-galactosidase (β-Gal). (E) The upd3-lacZ staining intensity was quantified in *N*=3 independent experiments. (F) *Drosophila* guts expressing 10×STAT92E-GFP, a reporter of JAK-STAT activation. (F) Quantification of 10×STAT92-GFP staining intensity was based on *N*=3 independent experiments. \**P* < 0.05; \*\**P* < 0.01; \*\*\**P* < 0.001; \*\*\*\**P* < 0.0001 by unpaired *t*-test (A, B, C, D, E) or Mann-Whitney U test (C, D, F); ns, nonsignificant. All error bars are the SEM.

Since cell death in the midgut is often compensated by proliferation [34], we next investigated mitotic activity by immunostaining for phospho-histone H3 (pH3). Loss of Rab21 resulted in a significant increase in the percentage of proliferating cells compared to controls (Figure 3C). Importantly, we were able to rescue the increased proliferation after Rab21 RNAi by overexpressing Rab21^deg^ in enterocytes (Figure 3D).

Upon stress, damaged enterocytes help trigger the compensatory proliferation of intestinal stem cells by secreting the inflammatory cytokine upd3, which activates JAK-STAT signaling in the intestinal epithelium [40,49]. We thus hypothesized that depletion of enterocyte Rab21 leads to non-cell autonomous compensatory proliferation through the upd3-JAK-STAT signaling axis. Using a upd3 reporter line, we observed higher levels of the reporter upon Rab21 depletion from enterocytes (Figure 3E). Consistent with this observation, the 10×STAT92E-GFP reporter revealed a dramatic increase in JAK-STAT signaling activity in Rab21-depleted guts (Figure 3F). These data support the idea that increased proliferation in Rab21-depleted guts is due to a non-cell autonomous mechanism.

### Non-cell autonomous proliferation in the Rab21-depleted gut is caused by cell death and activation of the transcriptional co-activator yki

To confirm that non-cell autonomous proliferation was responsible of the increased amount of pH3+ cells in guts with Rab21-depleted enterocytes, we co-depleted upd3 and Rab21 in enterocytes using inducible upd3 RNAi. Depleting enterocyte upd3 was sufficient to suppress the overproliferation phenotype observed after Rab21 knockdown (Figure 4A). Enterocyte-derived upd3 activates various pathways in the *Drosophila* adult midgut, and among these, the Hippo and JNK pathways are well documented [34,36,50–52]. We performed epistasis experiments using inducible RNAi of basket (bsk) and yki to assess the requirements for the JNK and Hippo pathways, respectively, in mediating increased upd3 expression upon Rab21 depletion. Only yki inhibition was sufficient to attenuate the compensatory proliferation induced by Rab21 depletion (Figure 4B).

**Figure 4.**
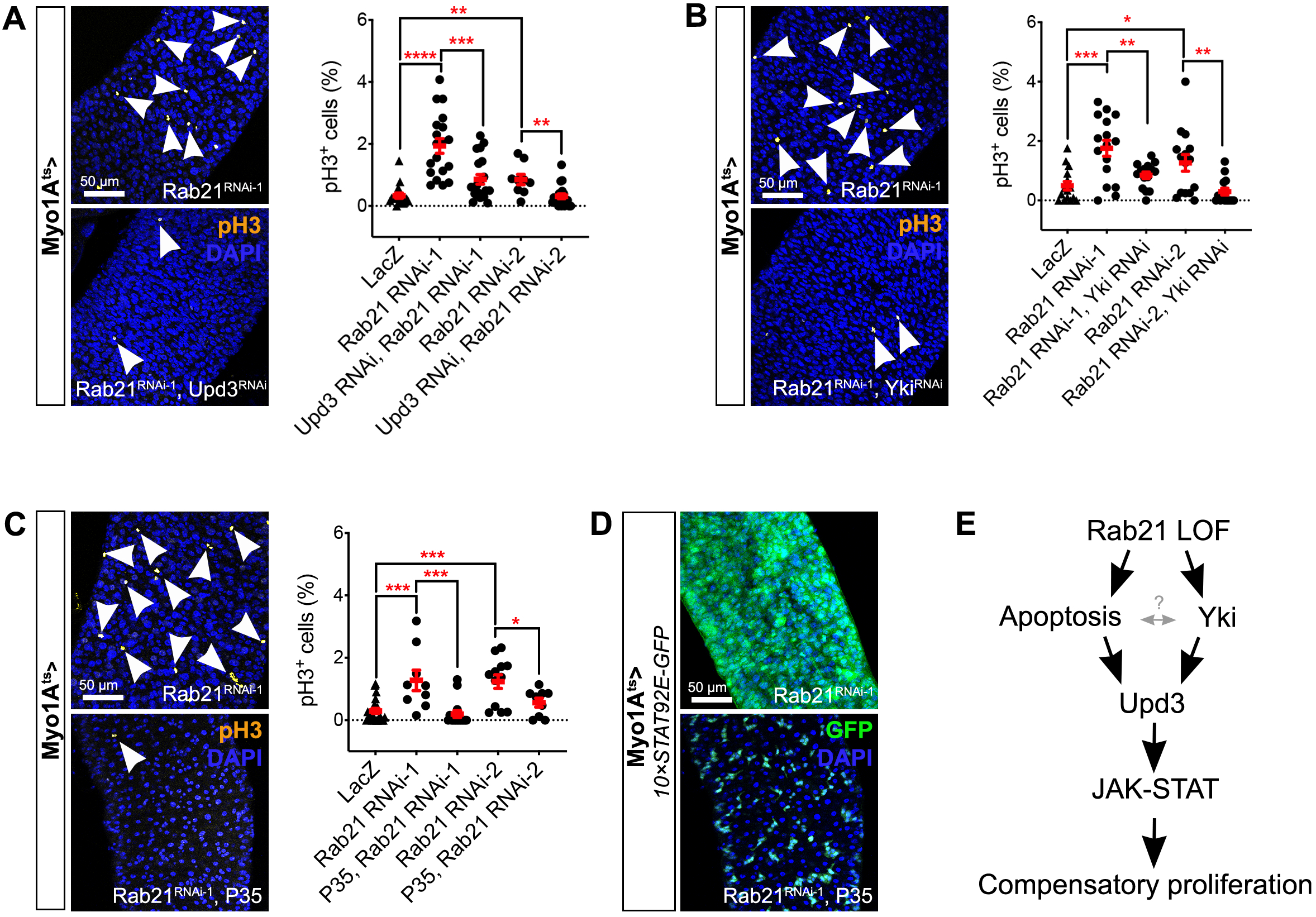
Rab21 restricts cell death and yki activation. (A) Mitotic cells were identified in guts with enterocytes depleted of Rab21 or Rab21/upd3 by immunostaining for pH3 (arrowheads). Quantification of pH3^+^ cells was based on *N*=3 independent experiments. (B) Mitotic cells were identified in guts with enterocytes depleted of Rab21 or Rab21/yki by immunostaining for pH3 (arrowheads). Quantification of pH3^+^ cells was based on *N*=3 independent experiments. (C) Guts with Rab21-depleted enterocytes with and without P35 expression were immunostained for pH3 (arrowheads). Quantification of pH3^+^ cells was based on *N*=3 independent experiments. (D) Expression of the 10×STAT92E-GFP reporter in *Drosophila* guts with Rab21-depleted enterocytes with and without P35 expression. Images are representative of *n*=16 guts in total from *N*=3 independent experiments. (E) Schematic of the signaling pathways affected by Rab21 loss-of-function in enterocytes. *\*P* < 0.05; \*\**P* < 0.01; \*\*\**P* < 0.001; \*\*\*\**P* < 0.0001 by unpaired *t*-test (A, B, C) or Mann-Whitney U test (A, B, C); ns, non-significant. All error bars are the SEM.

As we observed an increase in apoptotic cells in guts depleted of Rab21, we also evaluated whether cell death participates to the increased mitotic activity and upd3-related inflammatory phenotypes. To do so, we blocked apoptosis by overexpressing the baculovirus protein P35 in Rab21-depleted enterocytes. In this context, proliferation (Figure 4C) and JAK-STAT activation (Figure 4D) were both largely diminished compared to controls, demonstrating that cell death also increased upd3 secretion to mediate JAK-STAT activation. These data show that enterocyte Rab21 is required to block yki activation and enhance enterocyte survival, which both participate in restricting intestinal stem cell proliferation in a non-cell autonomous manner (Figure 4E).

### Rab21 is required for proper endolysosomal regulation in enterocytes

Rab GTPases are important regulators of cellular trafficking events [7], and Rab21 has been associated with diverse trafficking processes, mainly at early endosomes [20,22,44,45]. Therefore, to gain a better understanding of the dysfunctional enterocytic processes causing the physiological responses in the gut, we investigated early endosomal compartments upon Rab21 depletion. Using the GFP:2×FYVE probe, which labels phosphatidylinositol-3-phosphate (PtdIns(3)P) present at early endosomal membranes, we observed that the PtdIns(3)P pool was decreased in Rab21-depleted enterocytes compared to controls (Suppl. Fig. 4A). In addition, immunostaining for the early endosomal marker hepatocyte growth factor regulated tyrosine kinase substrate (Hrs), a well-known PtdIns(3)P-interacting protein [53], revealed decreased Hrs^+^ dots upon Rab21 knockdown compared to controls (Suppl. Fig. 4B). These data suggest that Rab21 regulates PtdIns(3)P more directly than previously thought [21]. Our previous work characterizing the RAB21 interactome [22] identified close proximity between RAB21 and the phosphatidylinositol 3-phosphate 5-kinase (phosphoinositide kinase, FYVE-type zinc finger containing) that is required to generate phosphatidylinositol 3,5-biphosphate (PtdIns(3,5)P2) from PtdIns(3)P. We therefore also assessed whether the decreased PtdIns(3)P level was due to excessive generation of PtdIns(3,5)P2. Using the mCherry:ML1N2× probe, which labels PtdIns(3,5)P2, we noticed a significant decrease in PtdIns(3,5)P2 pools in Rab21-depleted guts compared to controls (Suppl. Fig. 4C). PtdIns(3,5)P2 localizes to late endosomal and lysosomal membranes; therefore, we investigated whether these compartments were affected by Rab21 depletion. We observed no significant differences in Rab7^+^ vesicles (Suppl. Fig. 4D), suggesting that late endosomes were not affected. In addition, we used the GFP-lysosomal-associated membrane protein (LAMP) construct [54] to assess lysosome localization upon Rab21 knockdown. As observed in other fly tissues and in mammalian cells [22,44], we did not notice any differences in the lysosomes of control and Rab21-depleted enterocytes (Suppl. Fig. 4E). These data show that Rab21 depletion perturbs specific phosphoinositide pools, potentially impacting enterocyte sorting pathways. Moreover, as membrane trafficking and cellular signaling are closely related, the effects of enterocytic Rab21 depletion on endosomal trafficking may impact signaling pathways responsible for the observed phenotypes.

### Autophagy and EGFR signaling are independent in enterocytes

To gain deeper insight into the trafficking processes that regulate enterocyte functions, we performed a genetic screen to identify genes phenocopying the increased compensatory proliferation and inflammation caused by Rab21 depletion, by assaying pH3 and upd3 levels, respectively. We focused on endosomal genes linked to different membrane trafficking steps or processes (Figure 5A), as well as genes associated with functions ascribed to Rab21: autophagy regulation (Autophagy-related 1 (Atg)1, *Atg4,* Vesicle-associated membrane protein 8 (*Vamp8),* and Syntaxin 17 (*Syx17*)), Wiskott Aldrich Syndrome protein and scar homologue (WASH)/Retromer cargo regulation (washout (*wash),* Strumpellin (*Strump),* Family with sequence similarity 21 (*FAM21*), Vacuolar protein sorting (*Vps26, Vps29,* and *Vps35),* β1 integrin recycling (myospheroid (*mys*)), and EGFR degradation (*Ras^V12^).* Strikingly, disrupting enterocyte autophagy (*Syx17, Vamp8, Atg4*, *Atg1*), early endosomal tethering (class C core vacuole/endosome tethering (CORVET) complex genes) (*Vps8*, *Vps18*) and early and late endosomal regulation (Rab5 and Rab7, respectively) led to increased proliferation (Figure 5B, C) and upd3 expression (Figure 5D, E). As previously characterized [55], EGFR-MAPK signaling (Ras^V12^) also led to increased proliferation and upd3 expression (Figure 5A-E). Similar results were observed upon expression of a DN form of shibire (shi; Figure 5B-E), while depletion of Adaptor Protein complex 2, α subunit (AP-2) had no effect (Suppl. Fig. 5A-B). Finally, loss of the Retromer subunit Vps26 (Figure 5B-E) also phenocopied Rab21 depletion; however, depleting two other Retromer subunits (Vps29 and Vps35) did not (Suppl. Fig. 5A-B). None of the other tested genes phenocopied both the over proliferative and inflammatory phenotypes observed upon Rab21 depletion (Suppl. Fig. 5A-B). We conclude that early endosomal functions (Vps26, Rab5, Vps8, and Vps18) and autophagy (shi, Atg4, Vamp8, and Syx17) are required in enterocytes for proper gut homeostasis.

**Figure 5.**
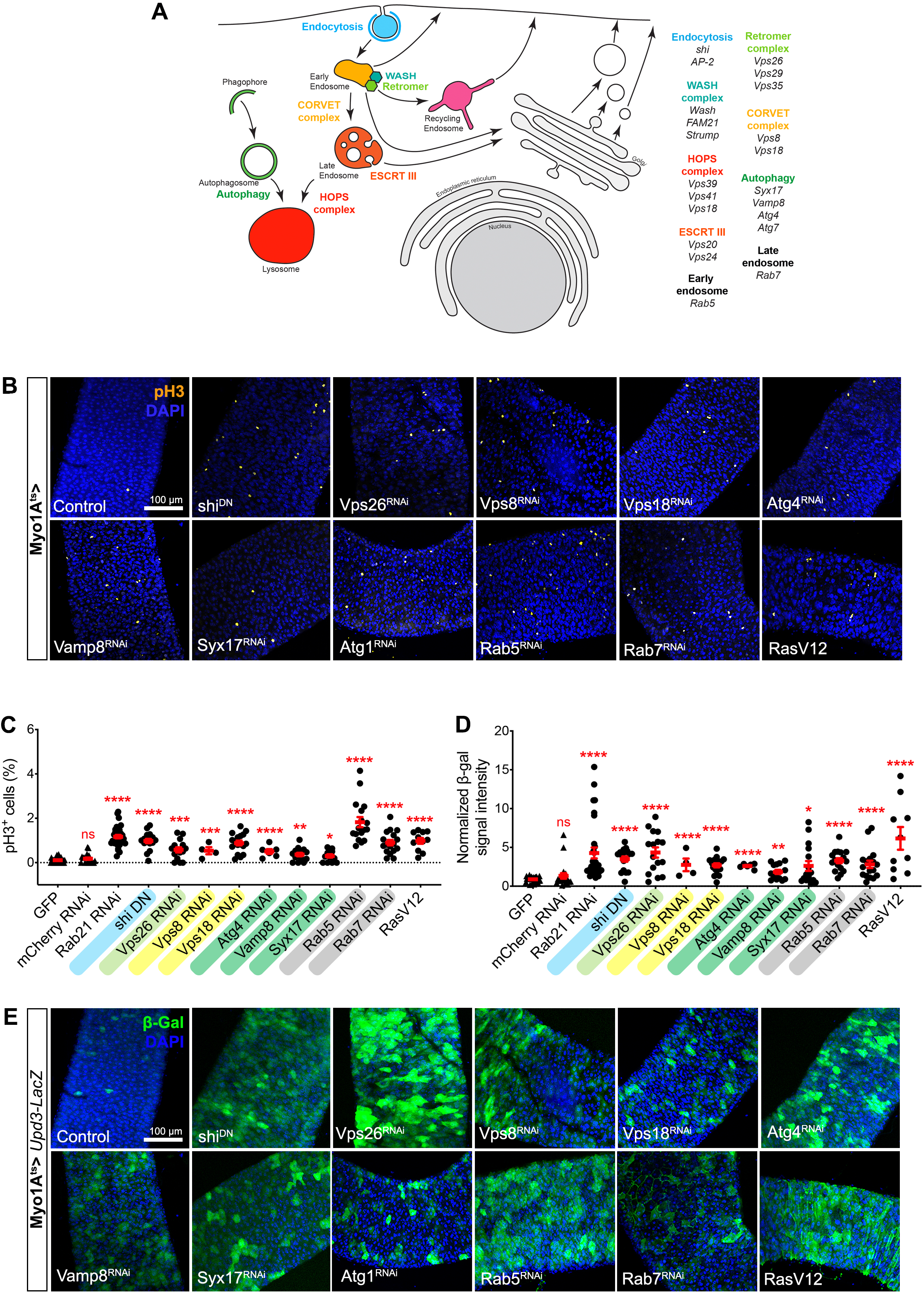
Depletion of autophagic genes phenocopies enterocyte Rab21 knockdown. (A) Schematic of the subcellular localizations of the genes assessed in the screen. (B) Mitotic cells were detected after depletion of Vps26, CORVET components, autophagy-related genes, Rab5, or Rab7, or after expression of shi^DN^ or constitutively active Ras, by immunostaining for pH3. (C) Quantification of pH3^+^ cells was based on *N*=3 independent experiments, except for Vps8^RNAi^ and Atg4^RNAi^ (*N*=1). Gene names are highlighted in colors corresponding to the membrane trafficking steps they affect (defined in A). (D-E) Expression of upd3-LacZ was detected after depletion of Vps26, CORVET components, autophagy-related genes, Rab5, or Rab7, or after expression of shi^DN^ or constitutively active Ras, by immunostaining for β-galactosidase (β-Gal). (D) The upd3-LacZ staining intensity was quantified based on *N*=3 independent experiments, except for Vps8^RNAi^ and Atg4^RNAi^ (*N*=1). Gene names are highlighted as in C. \**P* < 0.05; \*\**P* < 0.01; \*\*\**P* < 0.001; \*\*\*\**P* < 0.0001 by Mann-Whitney U test (C, D); ns, non-significant. All error bars are the SEM.

Activated Ras phenocopied Rab21 depletion (Figure 5B-E and [55]), and interestingly, recent work has highlighted a role for autophagy in regulating EGFR signaling in *Drosophila* intestinal stem cells [56,57]. Therefore, we assessed if EGFR signaling was impaired upon Rab21 knockdown by immunostaining for diphosphorylated extracellular signal-regulated kinase ((dp)ERK). Guts depleted of Rab21 displayed a significant increase in dpERK compared to controls (Figure 6A). Importantly, overexpression of a DN form of Egfr (Egfr^DN^) was sufficient to rescue the non-cell autonomous proliferation of intestinal stem cells induced by Rab21 knockdown (Figure 6B). These data demonstrate that Rab21 contributes to the regulation of enterocyte EGFR signaling. We next investigated if this contribution could be mediated by regulating autophagy. When we immunostained guts for refractory to sigma P (ref(2)p), an autophagic cargo protein, we noticed an increase in ref(2)p^+^ dots in Rab21-depleted guts compared to controls (Suppl. Fig. 6A). This suggested a deregulation of autophagic flux, as previously observed in other cell types [44,58]. These data were further supported by transmission electron microscopy (TEM), which revealed an increased number of autophagosomes, recognizable by their double membranes, upon Rab21 depletion (Suppl. Fig. 6B, arrowheads). To validate the involvement of Rab21’s autophagic function in the regulation of EGFR signaling, we assessed dpERK signal intensity after depleting autophagy-related genes. Surprisingly, no significant differences were observed (Figure 6C; Suppl Fig. 6C), with the exception of Syx17 RNAi (Figure 6C). In addition, inhibition of EGFR signaling was unable to rescue the increased mitotic activity observed in guts depleted of autophagic genes (Figure 6D). These data show that Rab21 contributes to both EGFR signaling and autophagy regulation; however, these effects are independent of each other. These results also highlight that contrary to intestinal stem cells, autophagy and EGFR signaling are not epistatic in enterocytes.

**Figure 6.**
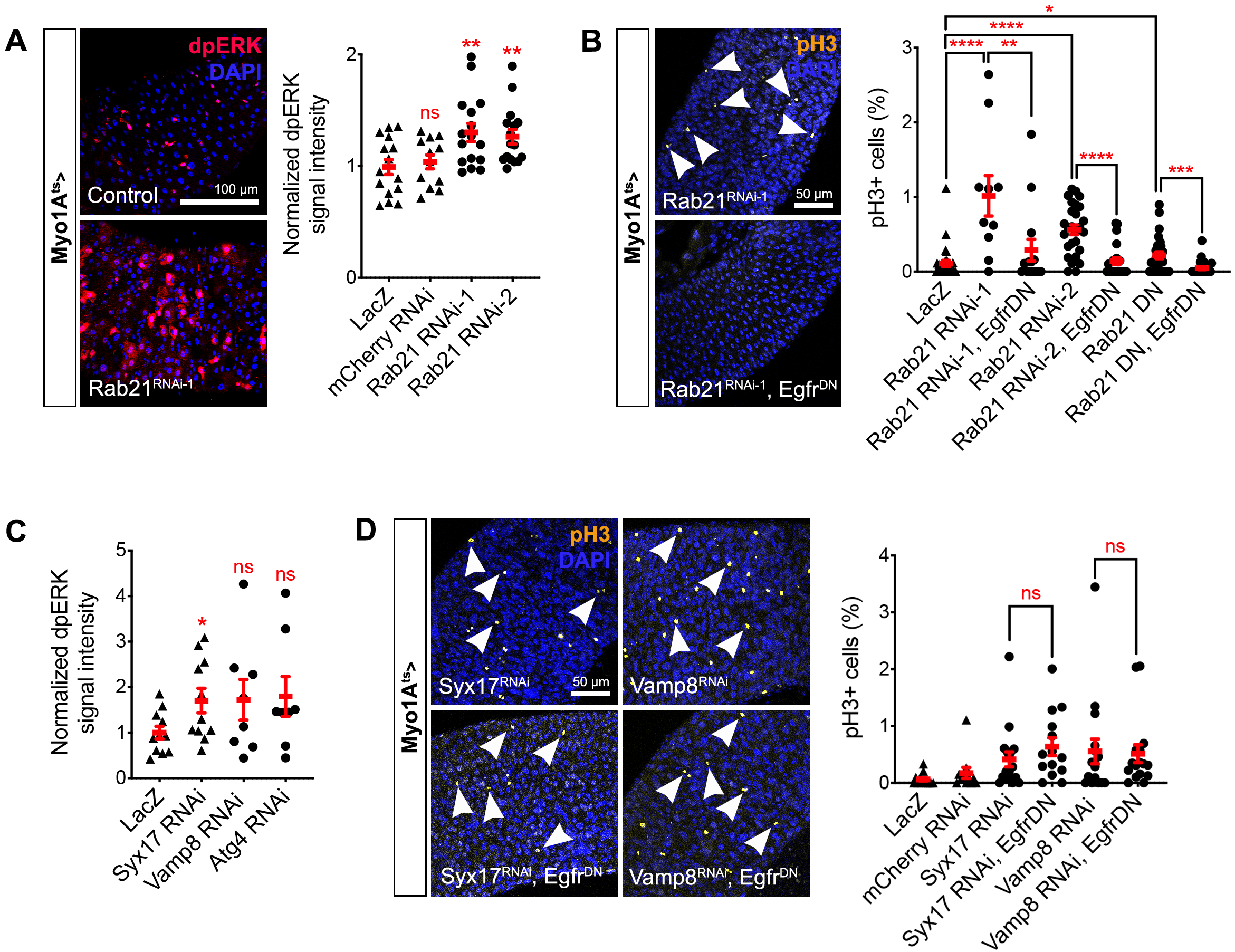
Rab21 regulates EGFR signaling in enterocytes independently of autophagy. (A) ERK activation was examined after Rab21 depletion by immunostaining for dpERK. (A) Quantification of dpERK signaling intensity was based on *N*=3 independent experiments. (B) Mitotic cells were detected after depleting Rab21 from enterocytes with and without Egfr^DN^ overexpression by immunostaining for pH3 (arrowheads). Quantification of pH3^+^ cells was based on *N*=3 independent experiments. (C) Quantification of dpERK signal intensity upon depletion of autophagic proteins in *N*=2 independent experiments. (D) Mitotic cells were detected after depletion of autophagic proteins from enterocytes with and without Egfr^DN^ overexpression by immunostaining for pH3 (arrowheads). Quantification of pH3^+^ cells was based on *N*=3 independent experiments. \**P* < 0.05; \*\**P* < 0.01; \*\*\**P* < 0.001; \*\*\*\**P* < 0.0001 by unpaired *t*-test (A, C) or Mann-Whitney U test (B, D); ns, non-significant. All error bars are the SEM.

### Enterocyte Rab21 knockdown affects solute carrier transporter abundance

The above data genetically identified the cellular signaling pathways affected by Rab21 depletion from enterocytes. However, they do not provide an unbiased view of the proteins affected by Rab21 knockdown. Therefore, to supplement our genetic analyses, we systematically identified proteins affected by Rab21 depletion in enterocytes. To do so, we performed a TMT-based quantitative proteomic analysis of control fly guts and those with enterocyte-specific Rab21 depletion in biological triplicate. We identified 2,691 proteins in total, of which 101 were differentially modulated more than two folds (Figure 7A), with 57 proteins that increased and 44 that decreased (Figure 7B) in Rab21-depleted cells compared to controls. Interestingly, many of the increased proteins belonged to the solute carrier (SLC) transporter family (Figure 7B, highlighted with a star). Previous studies have demonstrated functional links between Rab21 and certain SLC members [22,25], supporting our data. The proteomic approach also revealed decreased abundance of several proteins in the cytochrome P450 (CYP) family (Figure 7B, square), as well as proteins related to proteolysis (Figure 7B, circle). Finally, three proteins related to lipid metabolism were also deregulated upon Rab21 depletion from enterocytes (Figure 7B, hash symbol). We then investigated Reactome pathways enriched with differentially regulated proteins (Figure 7C). This analysis revealed the deregulation of several processes related to sugar: “lysosomal oligosaccharide catabolism” was decreased, while “cellular hexose transport”, “intestinal hexose absorption”, and “glucogenesis” were increased. Consistent with the increase in SLC proteins, “SLC-mediated transmembrane transport” was also enriched. Interestingly, some of the upregulated SLC proteins are involved in glucose transport, including proteins belonging to the SLC2, SLC5 and SLC16 families. From these data, we conclude that Rab21 is likely required for the proper regulation of SLC, CYP, and lipid metabolism proteins, and maintains proper absorption of sugar and potentially other nutrients by enterocytes.

**Figure 7.**
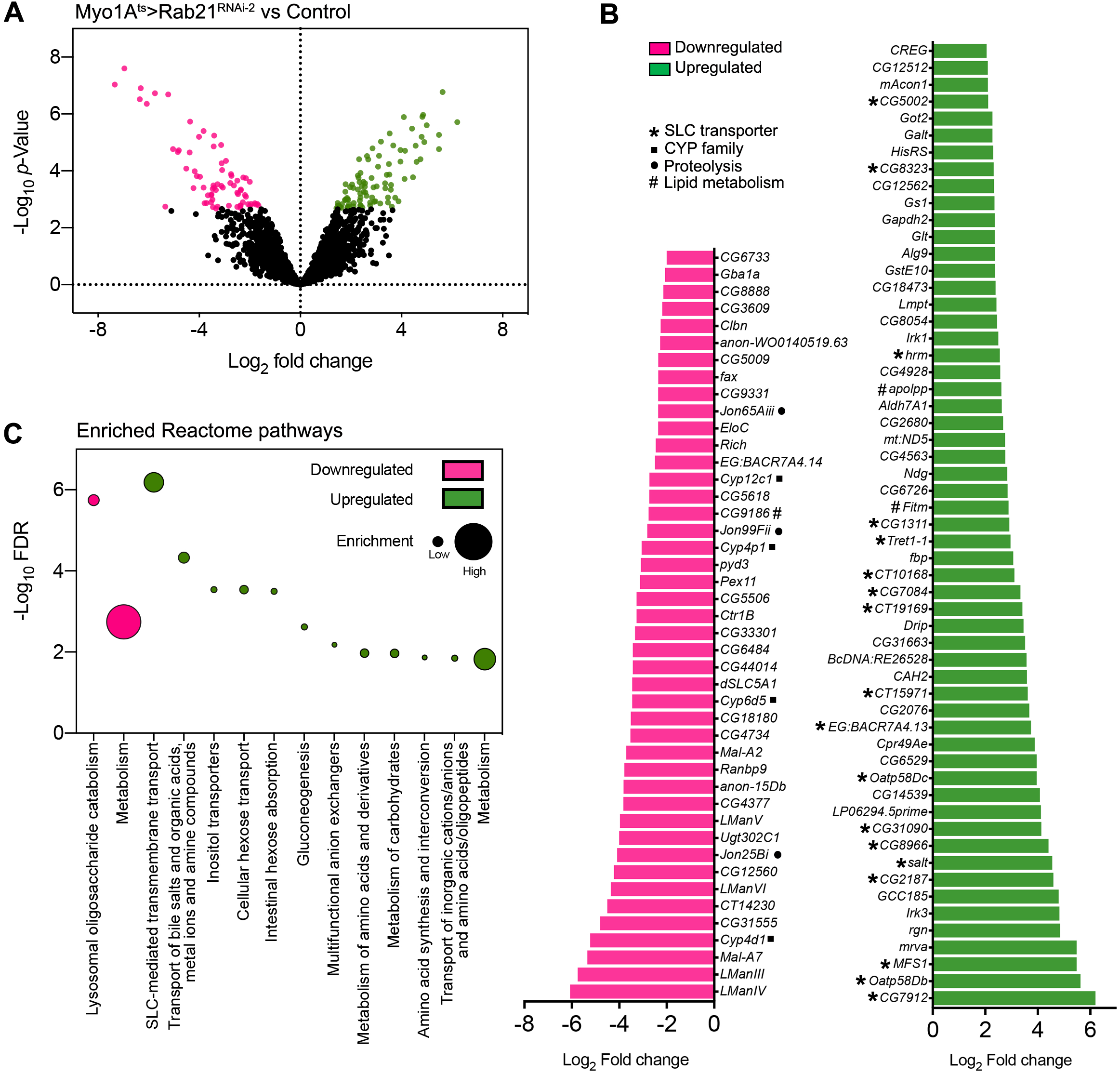
TMT-based quantitative proteomic analysis of *Rab21-depleted* guts. (A) Volcano plot of protein abundance differences between Myo1A^ts^>*Rab21*^RNAi-2^ and control guts in *N*=3 independent biological replicates. Pink indicates a FDR < 0.05 and a log2 fold change (FC) < −1.5; while green indicates a FDR < 0.05 and a log2 FC > 1.5. (B) Genes associated with proteins with significant (FDR < 0.05) decreases (pink) or increases (green) in abundance (log2 FC of < −2 and > 2, respectively), with their associated FCs. (C) Reactome pathways enriched with proteins with significantly (FDR < 0.05) decreased (pink) or increased (green) abundance (log2 FCs of < −2 and > 2, respectively). The distance of each dot from the horizontal axis represents the −log10 FDR of that pathway. The diameter of each dot corresponds to the number of proteins with modified abundance in that specific pathway compared to the total number of proteins with modified abundance.

## DISCUSSION

The importance of the membrane trafficking machinery in cell homeostasis is well established [2]; however, questions remain regarding the cell-specific functions of the majority of its components. In enterocytes, previous studies uncovered a role for endosomal recycling in coordinating apical-basal axis formation [4,8,9] and brush border formation [4,8]. Here, we found that the early endosomal protein Rab21 is required in enterocytes to maintain tissue homeostasis through the regulation of multiple signaling events crucial for their survival and function.

We showed that Rab21 knockdown in enterocytes leads to the formation of a multilayered epithelium with aberrant localization of the adherens and septate junction markers arm and cora. Intriguingly, while flies depleted of enterocyte Rab21 have shorter lifespans, epithelium integrity is conserved, indicating that the reduced lifespan is not due to gut leakiness. Similarly, flies depleted of intestinal septate and adherens junction components (Tetraspanin 2A and E-Cadherin, respectively) do not show tissue permeability defects [59,60]. Therefore, cora and arm mislocalization in Rab21-depleted guts may reflect defects in tissue homeostasis and related signaling events rather than impairment of junction formation and function.

Our data clearly illustrate that cellular homeostasis is dramatically perturbed upon Rab21 depletion from enterocytes. Indeed, the enterocyte population diminished while the intestinal stem cell and enteroendocrine cell populations expanded. The results indicate that the reduction in enterocytes is due to induction of apoptosis, and that compensatory proliferation is responsible for the rise in intestinal stem cells. However, it is unclear why we observed expansion of the enteroendocrine cell population. Previous studies highlighted a role for the JAK-STAT pathway in enteroendocrine cell specification [61,62]. A high level of JAK-STAT signaling is required for enteroendocrine fate specification, demonstrated by the fact that the intestinal epithelia of hypomorphic *Stat92^E06346^* mutant flies are composed mainly of enterocytes [61]. Interestingly, we showed that guts depleted of Rab21 display a massive increase in JAK-STAT signaling activity. Therefore, it is conceivable that the increase in enteroendocrine cells is caused by hyperactivity of the JAK-STAT pathway.

In addition, we observed the induction of inflammation upon Rab21 depletion. This inflammation was mediated by upd3 secretion and led to compensatory proliferation. Enterocytes secrete inflammatory cytokines in response to a large range of stresses (e.g., apoptosis, infection, reactive oxygen species, JNK activation, injury) [34–36,38,39,63,64], with upd3 being the most abundant [34]. Secretion of upd3 triggers compensatory proliferation by activating JAK-STAT signaling in intestinal stem cells and visceral muscle cells [34,39,61]. Visceral muscle cells, in turn, secrete the Egfr ligand vein, which promotes intestinal stem cell proliferation through Egfr [39,65]. Various signaling pathways act as stress sensors in the intestinal epithelium, including the Hippo-yki, JNK, and p38 pathways [34–36,38,63]. In enterocytes, activation of any one of these pathways stimulates upd3 secretion [34–36,38,63]. Interestingly, constitutive activation of the JNK pathway induces cell death [34]. However, enterocyte apoptosis is not responsible for JNK-mediated compensatory proliferation, since JNK-induced compensatory proliferation cannot be rescued by P35 expression [34]. In addition to upd3 secretion, stressed enterocytes participate in promoting intestinal stem cell proliferation through the transcriptional activation of the Egfr ligand maturation factor rhomboid, which promotes the secretion of mitogenic Egfr ligands [66]. Consistent with these data, in Rab21-depleted guts, we were not able to rescue increased proliferation by inhibiting JNK signaling, while blocking apoptosis was sufficient to significantly reduce compensatory proliferation. Furthermore, we showed that yki was required for the compensatory proliferation of intestinal stem cells after Rab21 depletion. The Hippo-yki pathway plays well-characterized roles in promoting cell growth/proliferation and inhibiting cell death [67]; therefore, the relationship between yki signaling and apoptotic signaling in triggering compensatory proliferation in Rab21-depleted guts remains unclear. It is likely that these pathways act synergistically to mediate the non-cell autonomous proliferation of intestinal stem cells upon Rab21 knockdown, and may not be individually sufficient for this purpose. In accordance with this hypothesis, recent work showed that in unchallenged conditions, yki depletion from enterocytes tended to diminish enterocyte apoptosis, while hippo inhibition tended to increase it [37]. Such data suggest context-dependent requirements for yki functions, with the Hippo-yki pathway involved in enterocyte apoptosis in normal conditions.

The compensatory proliferation and inflammation phenotypes associated with Rab21 depletion from enterocytes are similar to those observed in Rab11-depleted guts [9,12]. However, these phenotypes are related to distinct signaling dysfunctions. We showed that Rab21, in enterocytes, negatively regulates EGFR signaling, while Rab11 functions independently of this pathway [9,12]. Overexpression of Egfr^DN^ in Rab21-depleted guts suppressed compensatory proliferation. Consistent with these findings, in HeLa and HEK293T cells, RAB21 is required for proper EGFR internalization and degradation [45], although the specific molecular mechanism by which it regulates EGFR remains to be defined. Our genetic screen revealed that depletion of autophagy- and endosome-linked genes from enterocytes phenocopied knockdown of Rab21. Furthermore, constitutive activation of EGFR-MAPK signaling led to similar results, as previously observed [55]. Recently, an autophagy-endocytosis network was linked to the negative regulation of EGFR signaling in intestinal stem cells [56,57]. However, EGFR pathway regulation appears to be different in enterocytes, as inhibiting EGFR signaling did not rescue autophagy-induced increases in proliferation. Therefore, our data show that autophagy and EGFR signaling are independent in enterocytes, yet are both regulated by the early endosomal protein Rab21.

We showed that Rab21 depletion negatively affects PtdIns(3)P and PtdIns(3,5)P2 pools, as well as Hrs^+^ early endosomes. We hypothesize that these defects result in impaired endosomal trafficking and inappropriate Egfr trafficking. Therefore, the increased Egfr activity could result from decreased Hrs endosomal recruitment or inefficient endosomal maturation. We did observe normal late endosomes and lysosomes, suggesting defects in earlier trafficking stages. This was consistent with the phenotypes of guts containing enterocytes depleted of Rab5 and CORVET complex components.

Proper regulation of EGFR signaling in early enterocytes is crucial for their morphology and maturation [65,68]. Previous studies identified that EGFR-MAPK activation in enterocytes induces their endoreplication [68] and that overexpression of Egfr triggers enterocyte delamination [65]. Interestingly, joint activation of Egfr and mir-8 stem loop in the wing disc epithelia leads to polyploid cells that induce apoptosis in neighboring cells [69]. From these observations, we hypothesize that Rab21 depletion from enterocytes might lead to EGFR-MAPK activation in these cells, which would in turn trigger the death of surrounding cells, with both effects required for the observed Rab21-related phenotypes.

Finally, to uncover the physiological functions altered by defective early endosomal trafficking, we performed a TMT-based quantitative proteomic analysis. The elevated levels of multiple SLC families are consistent with previously reported roles for Rab21 [22,23], and suggests that SLC transporters accumulate either intracellularly or at the plasma membrane. Rab21 has recently reported to be involved in clathrin- and dynamin-independent endocytosis [22,23]. Thus, both scenarios are consistent with Rab21 functions.

However, we favor the possibility that SLCs accumulate in intracellular vesicles that are improperly targeted for degradation, or that they are not recycled efficiently to the plasma membrane and are present in aberrant/inefficient recycling endosomes, given that Rab21 depletion sensitizes enterocytes to cell death. We posit that the opposite would occur in conditions where more transporters would be at the cell surface. Intriguingly, the SCL2 (CG8837), and SLC5 (CG2187, salt, CG8966, CG31090) families were the most highly represented, and both are involved in sugar transport [28,70,71], consistent with enrichment in sugar-related Reactome pathways. In the future, it will be interesting to determine if loss of enterocyte sugar import contributes to their death, and to identify the responsible SLC(s). Our data also reveal decreases in several proteins related to proteolysis, which are likely digestive enzymes. Prior to absorption, digestive enzymes are responsible for breaking down ingested macromolecules before their absorption by enterocytes [3]. Therefore, decreased digestive enzymes might result in improper absorption, which could also lead to dysfunctional enterocytes and decreased lifespan. Finally, we demonstrated decreased abundancies of CYP family members [72,73]. CYPs are a superfamily of enzymes with heme and iron binding functions as well as oxidoreductase activity [74]. Decreased amounts of these enzymes in enterocytes could lead to decreased detoxification capacities in these cells, which could also account for their death. Untangling these observations will be of great interest in our future studies.

To conclude, our data shed light on the importance of the enterocyte early endosomal machinery in maintaining proper tissue homeostasis. Our results demonstrate that in enterocytes, Rab21 acts differently than Rabs previously investigated in this cell type. We show that Rab21 regulates the EGFR pathway and autophagy, although independently of each other. Moreover, we identify deregulation of specific SLC families, digestive enzymes, and CYP proteins, indicating physiological defects in specific cellular processes. Further investigations of the cell-specific functions of membrane trafficking regulators will highlight their underappreciated roles in tissue and organismal homeostasis.

## MATERIALS AND METHODS

### *Drosophila* strains

Fly stocks were maintained at 25°C on a standard diet composed of 7 g/L agar, 60 g/L cornmeal, 60 g/L molasses, 23.5 g/L yeast extract, 4.5 mL/L (BioShop), and 4 mL/L propionic acid. Genotypes used in this study were: Rab21-GAL4 (*BDSC_51593),* UAS-GFP:Rab21 WT [21], UAS-GFP:Rab21^T27N^ DN [21], UAS-GFP:Rab21^Q73L^ CA [21], Myo1A-GAL4; Tub-GAL80ts, UAS-GFP [49], UAS-LacZ (*BDSC_1776*) UAS-mCherry RNAi (*BDSC_35785),* UAS-Rab21 RNAi-1 (*VDRC_32941),* UAS-Rab21 RNAi-2 (*VDRC_109991),* Dl:eGFP (*BDSC_59819),* Myo1A-GAL4, Tub-GAL80ts; upd3-LacZ [39], UAS-10×STAT92E-GFP (*BDSC_26197),* UAS-P35 (*BDSC_5073),* UAS-upd3 RNAi (*BDSC_32859),* UAS-yki RNAi (*BDSC_34067),* UAS-GFP:2×FYVE [75], UAS-mCherry:ML1N2, UAS-mys RNAi (*BDSC_27735),* UAS-shi^K44A^ DN (*BDSC_5811),* UAS-Vps26 RNAi (*BDSC_38937),* UAS-Vps8 RNAi (*VDRC_ 105952),* UAS-Vps18 RNAi (*BDSC_54460),* UAS-Atg4 RNAi (*BDSC_35740),* UAS-Vamp8 RNAi (NIG-FLY), UAS-Syx17 RNAi (*BDSC_25896),* UAS-Atg1 RNAi (*BDSC_26731),* UAS-Rab5 RNAi (*VDRC_34096),* UAS-Rab7 RNAi (*VDRC_40337),* UAS-Ras^V12^ (*BDSC_64196*). UAS-AP-2 RNAi (*BDSC*_32866), UAS-Egfr DN (*BDSC_3364),* UAS-wash RNAi (*BDSC_62866),* UAS-Strump RNAi (*BDSC_51906),* UAS-FAM21 RNAi (*BDSC_50571),* UAS-Vps35 RNAi (*VDRC_ 45570),* UAS-Vps39 RNAi (*VDRC_40427),* UAS-Vps29 RNAi (*BDSC_53951),* UAS-Vps20 RNAi (*BDSC_40894*) UAS-Vps24 RNAi (*BDSC_38281),* and UAS-Vps41 RNAi (*BDSC_34871).*

New genotypes generated in this study were: (1) UAS-Rab21^deg^; (2) Myo1A-GAL4, Tub-GAL80ts; UAS-Rab21^deg^; (3) Myo1A-GAL4, Tub-GAL80ts; Dl:eGFP; (4) UAS-Rab21 RNAi-1, UAS-10×STAT92E-GFP; (5) UAS-10×STAT92E-GFP; UAS-LacZ; (6) Myo1A-GAL4, Tub-GAL80ts; UAS-P35; (7) Myo1A-GAL4, Tub-GAL80ts; UAS-upd3 RNAi; (8) Myo1A-GAL4, Tub-GAL80ts; UAS-yki RNAi; (9) Myo1A-GAL4, Tub-GAL80ts; UAS-EgfrDN; (10) Myo1A-GAL4, Tub-GAL80ts; UAS-mCherry:ML1N2; and (11) Myo1A-GAL4, Tub-GAL80ts; UAS-GFP:2×FYVE.

### Generation of the UAS-Rab21^deg^ transgenic line

The *Drosophila* Rab21 coding sequence was modified to prevent RNAi binding without affecting the amino acid sequence by modifying the third nucleotide of each codon present in the sequences targeted by Rab21 RNAi-1 and 2. The Rab21 degenerated coding sequence was synthetized by Integrated DNA Technology and cloned into the pUASt-AttB plasmid to generate pUASt-dRab21^deg^. Transgenic flies were generated by Genome Prolab through phiC31 transgenesis at the attP2 landing site.

### Immunofluorescence

*Drosophila* guts were dissected in 1× phosphate-buffered saline (PBS) over 20 minutes, fixed for 2 hours in 4% paraformaldehyde, and rinsed three times in PBS. To allow food to exit the lumen, the guts were incubated for 20 minutes in 50% glycerol/PBS and 10 minutes in PBS-0.1% Triton X-100 (PBT). Intestines were then blocked for 1 hour in 20% goat serum/PBT and processed for immunostaining. Primary antibody incubations were performed overnight at 4°C in PBT. The following primary antibodies were used: mouse anti-cora (C566.9, Developmental Studies Hybridoma Bank (DSHB), Iowa City, IA, USA; 1/100), mouse anti-arm (N2 7A1, DSHB; 1/100), mouse anti-pros (MR1A, DSHB; 1/100), mouse anti-Dl (C594.9B, DSHB; 1/100), mouse anti-β-galactosidase (Promega #Z3781; 1/500), rabbit anti-pH3 (Ser10; #06-570, Millipore, Oakville, ON, Canada; 1/1000), rabbit anti-cleaved caspase 3 (#9661, Cell Signaling Technology, Danvers, MA, USA; 1/500), rabbit anti-ref(2)p (ab178440, Abcam; 1/500), rabbit anti-dpERK (Thr202/Tyr204, #4370, Cell Signaling Technology; 1/500), and mouse anti-Hrs (Hrs 27-4, DSHB; 1/100). Fluorescent secondary antibody incubation was performed for 2 hours at room temperature (RT) in PBT. The following secondary antibodies were used (all from Thermo Fisher Scientific): anti-mouse Alexa Fluor 546 (#A11003; 1/500), anti-mouse Alexa Fluor 488 (#A11029; 1/500), anti-rabbit Alexa Fluor 546 (#A11035; 1/500), and anti-rabbit Alexa Fluor 488 (#A11008; 1/500). Phalloidin-Alexa Fluor 546 (#A22283, Thermo Fisher Scientific, 1/500) staining was performed at the same time as secondary antibody incubation. Tissues were mounted in SlowFade Gold Antifade Mountant with DAPI (#S36938, Thermo Fisher Scientific).

### *In situ* hybridization

Day 1: Intestines were fixed in 4% formaldehyde for 45 minutes at RT, rinsed three times in PBT, and dehydrated using an ethanol gradient in PBT (25%, 50%, 75% and 100% ethanol for 5 minutes each at RT). Before hybridization, tissues were successively rehydrated in 50% methanol/PBT and PBT for 10 minutes each at RT. Intestines were transferred to 50% hybridization buffer (HB; 50% deionized formamide, 5× Saline Sodium Citrate (SSC), 0.1% Triton X-1000, 2 mM vanadyl ribonucleoside, 100 μg/mL sonicated salmon sperm DNA, 100 μg/mL tRNA, and 100 μg/mL bovine serum albumin (BSA)) in PBT, then to 100% HB. Finally, guts were transferred to warm HB for 2 hours at 56°C, then incubated in 100 μL of HB containing 200 ng of digoxigenin (DIG)-labelled RNA probe overnight at 56°C. Day 2: Tissues were successively washed in warm 75%, 50%, 25%, and 0% HB in PBT for 15 minutes at 56°C. Endogenous peroxidases were quenched using 1% H2O2, and tissues were blocked in PBT containing 100 μg/mL BSA for 10 minutes at RT before incubation with horseradish peroxidase (HRP)-conjugated anti-DIG antibody (1/400) for 2 hours at RT. Intestines were washed five times in PBS for 5 minutes each at RT. A TSA Cyanine 3 Signal Amplification System (#SAT704A001EA, Perkin Elmer) was used for signal detection according to the manufacturer’s instructions. The *Drosophila* Rab21 RNA probe was produced from a reverse transcription-polymerase chain reaction (RT-PCR) library generated from adult *Drosophila*, using the following primers: Fwd-5’-AGATGAATTCTCCCTGGAGGATGGGAGAAG-3’, Rev-5’-GCTAGAATTCGTATCCGG ATTCTGGAGGC-3’.

### SYTOX staining

SYTOX Orange Nucleic Acid Stain (#S11368, Thermo Fisher Scientific) was diluted to 1 μM in 5% sucrose and fed to the flies overnight. Fly guts were then dissected in 1× PBS, incubated for 10 minutes in cold 1× PBS containing Hoechst, and were briefly rinsed in 1× PBS. Guts were mounted in SlowFade Gold Antifade Mountant with DAPI and imaged immediately.

### Confocal microscopy

Confocal images of guts were acquired on a Zeiss LSM 880 confocal microscope, using either a 20× Plan-APOCHROMAT/0.8 numerical aperture (NA) or a 40× oil Plan APOCHROMAT/1.4 NA objective. Confocal images represent maximum projections. Settings on the microscope were first adjusted on a control gut and maintained for the acquisition of the different conditions in each experiment. Images were analyzed in Fiji [76] or CellProfiler [77], then linearly thresholded and assembled into figure panels in Photoshop Version 21.1.1 (Adobe Systems, Inc., San Jose, CA, USA). All adjustments to contrast and other aspects of the images were performed similarly for all conditions in each experiment.

### TEM

Guts were dissected in 1× PBS, fixed in 2% glutaraldehyde in 0.1 M cacodylate buffer (pH 7.4), and stored at 4°C until processing [58]. Images were acquired on a Hitachi H-7500 transmission electron microscope. Images were analyzed in Fiji and thresholded in Photoshop.

### Gut RNA extraction and quantitative (q)RT-PCR

Guts (15–20) were quickly dissected in cold 1× PBS and transferred to TRIzol Reagent (#15596026, Invitrogen). RNA extractions were performed as described in the TRIzol Reagent protocol. RNA (500 ng) was reverse transcribed using the Maxima First Strand cDNA Synthesis kit (#K1671, Thermo Fisher Scientific). Primer sequences used for qRT-PCR were: hid Fwd-5’-CCCAGCCAACGATTTTTATG-3’, hid Rev-5’-TGTTACCGCTCCGGCTAC-3’, Actin Fwd-5’-CTCGCCACTTGCGTTTACAGT-3’, Actin Rev-5’-TCCATATCGTCCCAGTTGGTC-3’. Relative mRNA levels were calculated using the 2^-ΔΔCt^ method [78]. ΔCt values were obtained by normalizing to actin.

### Gut protein extraction

Guts (15-20) were quickly dissected in cold 1× PBS and transferred into 1× sodium dodecyl sulfate (SDS) loading buffer (50 mM Tris pH 6.8, 2% SDS, 250 μM dithiothreitol (DTT), 7.5% glycerol, 37.5 mM Tris pH 7.5, 112.5 mM NaCl, 750 μM ethylenediaminetetraacetic acid, and 0.75% Triton X-100) containing 1 × protease inhibitor (#P8340, Sigma-Aldrich). Guts were dissociated using a syringe and boiled for 5 minutes at 95°C. Protein extracts were incubated 15 minutes on ice, then centrifuged at 16,000 g for 15 minutes at 4°C. Supernatants were then transferred into new tubes and protein amounts were quantified using a BCA assay (#23225, Pierce). For immunoblots, protein extracts were resolved by 12% SDS-polyacrylamide gel electrophoresis and transferred to polyvinylidene difluoride membranes (#10600021, GE Healthcare).

### Survival curves

Flies were collected 1–3 days after eclosion and aged until death. Dead flies were scored every 2–3 days, and food was exchanged every other day.

### Smurf assays

Flies (1–3 days old) were collected in a tube and incubated at 29°C for 20 days. They were then transferred overnight to food containing FD&C blue dye #1 (1.75 g/100 mL). The next day, blue flies were scored as Smurf ^+^.

### Statistical analysis

Unpaired two-tailed Student’s *t*-tests and Mann-Whitney U tests were used to analyze significant differences between pairs of conditions. The proper statistical test for each experiment was determined by assessing the normal distributions of each condition in the experiment. Conditions with normal distributions were analyzed using unpaired two-tailed Student’s *t*-test, while conditions without normal distributions were analyzed using Mann-Whitney U tests. All statistical comparisons were performed using data collected from at least three biological replicates, unless otherwise specified. GraphPad Prism (GraphPad Software, La Jolla, CA, USA) was used for statistical analyses and graph generation.

### Protein processing for TMT-based proteomic analysis

Gut proteins were extracted as described above, using the same lysis buffer. Following gut dissection and protein extraction, biological replicates were flash frozen and simultaneously processed for TMT labelling. Proteins (50 μg) were reduced in 10 mM DTT for 2 minutes, then boiled and alkylated in 50 mM iodoacetamide for 30 minutes in the dark. DTT- and β-mercaptoethanol-free SDS loading buffer was added to the protein extracts to a final concentration of 1×. Proteins were resolved on 4–15% Mini-PROTEAN TGX Precast Protein Gels (#4561084, Bio-Rad) for 15 minutes at 200 V and visualized using SimplyBlue SafeStain solution (#LC6060, Thermo Fisher Scientific). For each condition, two bands (A and B) containing all stained proteins were excised from the gel. Gel bands were successively washed at RT with 300 μL of H2O and 300 μL of 100% acetonitrile (ACN) for 15 minutes each. The supernatants were removed, and the gel bands were successively washed in 300 μL of 50 mM triethylammonium bicarbonate (TEAB) and 300 μL of 50:50 ACN:50 mM TEAB at RT for 15 minutes each. Supernatants were discarded, and the previous last two washes were repeated if the gel bands remained blue. Finally, the gel bands were incubated in 150 μL of 100% ACN for 5 minutes at RT and dried for 5 minutes in a SpeedVac Vacuum Concentrator (Thermo Fisher Scientific). Ingel digestion was performed overnight in 50 μL reactions containing 12.5 ng/μL trypsin in 50 mM TEAB at 30°C. Digested gel bands were incubated in 50 μL ACN for 30 minutes at RT. Supernatants containing digested peptides were collected and residual peptides were eluted with 100% ACN and 1% formic acid. Peptides were dried and resuspended in 50 mM TEAB, and their concentrations were measured with a NanoDrop spectrophotometer (Thermo Fisher Scientific) using their absorbance at 205 nm. Finally, for each condition, the same maximal quantity of peptides was used for TMT labelling using a TMT10plex™ Isobaric Label Reagent Set (#90110, Thermo Fisher Scientific) according to the manufacturer’s protocol. Peptides from bands A and B from a same condition were independently labelled with the same label. Labelled peptides from all band A samples from the different conditions were then pooled, as were the labelled peptides from all band B samples. The pooled samples were dried, resuspended in 0.1% trifluoroacetic acid diluted in water, and desalted using a ZipTip. For each pooled sample, 1.5 μg peptides were resuspended in 1% formic acid diluted in water and loaded on a Q Exactive Orbitrap mass spectrometer (Thermo Fisher Scientific). Settings used were as previously described [22], except that the full-scan mass spectrometry (MS) survey spectra acquisition (*m/z* 375–1,400) was realized using a resolution of 140,000 with 3,000,000 ions and a maximum injection time of 120 ms.

### MS data analysis

For the TMT-based quantitative proteomic experiment, three independent biological replicates were used. Proteins were identified by MaxQuant using the UniProt *Drosophila melanogaster* proteome UP000000803. The “proteinGroup.txt” output file from the MaxQuant analysis [79] (Supplementary table 1) was used to collect corrected reporter intensities per sample for every detected protein, as a measure of their quantities in each sample. Since series of replicates were run on different days, we checked for the presence of batch effects by generating a multidimensional scaling plot using the limma v3.42.2 package [80] in the R environment. Once confirmed, batch effects were handled in R using the internal reference scaling (IRS) methodology [81], which is capable of correcting the random MS2 sampling that occurs between TMT experiments. Since the dataset contained proteins which were not quantified in all replicates, we first filtered for proteins that were identified in 2/3 replicates of at least one condition, and then we checked the spectra for missing data using the DEP v1.8.0 R package [82]. Once we verified that missing data were occurring at random, we imputed them using the k-nearest neighbor approach in DEP v1.8.0. Following this analysis, only proteins with 2 unique peptides were conserved for data representation (Figure 7B) and reactome enrichments (Figure 7C). Differential expression analysis was performed in R using DEP v1.8.0, with a false discovery rate (FDR) of 0.05 and a log2 fold change (FC) of 1.5. Enriched Reactome pathways were independently searched for proteins with >2-fold increases or decreases in abundance using the “STRING enrichment” plugin in Cytoscape [83]. The mass spectrometry data have been deposited to the ProteomeXchange Consortium via the PRIDE repository with the data set identifier PXD022413.

## ACKNOWLEDGMENTS

We thank M. Avino for helpful statistical analysis of the TMT experiment. We thank Bruce Edgard for kindly providing the transgenic fly lines Myo1-Gal4; Tub-Gal80^ts^,GFP and Myo1-Gal4,Tub-Gal80^ts^; and upd3-LacZ. We thank Nicole St. Denis for stylistic and copyediting services. We thank the proteomics platform at the Université de Sherbrooke for proteomic services, the histology and electronic microscopy platform for sample preparation, and the Photonic microscopy platform for use of a confocal microscope. We also thank all members of the Jean laboratory for relevant suggestions during the course of this work. Steve Jean and François-Michel Boisvert are members of the Fonds de Recherche du Québec-Santé (FRQS)-Funded Centre de Recherche du CHUS. Steve Jean is a recipient of a Research Chair from the *Centre de recherche médicale de l’Université de Sherbrooke.* Sonya Nassari is the recipient of a postdoctoral fellowship from the FRQS. This research was supported by operating grants from the Canadian Institutes of Health Research (CIHR) and by junior faculty salary awards from CIHR and FRQS to S.J.

## AUTHOR CONTRIBUTIONS

SN performed the experiments. DL and FMB assisted with all MS analyses. SJ and SN designed the experiments. SJ and SN wrote the manuscript.

