## SupplementaryFigures for "The early endosomal protein Rab21 is critical for enterocyte functions and intestinal homeostasis"

**Supplementary Figure legends**

**Supplementary Figure 1. Rab21 expression in the drosophila gut epithelium** (A) Phalloidin staining (to detect actin) in flies expressing GFP:Rab21wt under the control of the *Rab21* promoter. *N*=3 independent experiments. (B) Fluorescent *in situ* hybridization of Rab21 in $W^{1118}$ *Drosophila* gut. Arrowheads indicate polyploid cells. *N*=2 independent experiments. (C) GFP:Rab21wt-expressing guts were immunostained for prospero (pros) and Delta (Dl). Arrowheads indicate GFP:Rab21$^{+}$ pros$^{+}$ and GFP:Rab21$^{+}$ Dl$^{+}$ cells. *N*=3 independent experiments.

**Supplementary Figure 2. Loss of enterocyte Rab21 affects survival and gut cell density** (A) Inversed binary images of *Drosophila* guts stained with DAPI. (A) Cell density was quantified based on *N*=3 independent experiments. (B) Inversed binary images of Drosophila guts immunostained for armadillo (arm). Images are representative of *N*=3 independent experiments. (C) TEM of *Drosophila* gut tissue sections, highlighting enterocyte microvilli in control and Rab21-depleted enterocytes. (D) Smurf assays of intestinal permeability in control and Rab21-depleted flies. Rab21$^{RNAi}$, *n*=27; controls, *n*=16, from *N*=3 independent experiments. (E) Survival curves in control and Rab21-depleted enterocytes. *N*=3 independent experiments, except for Rab21$^{DN}$ (*N*=2). ****$P <$ 0.0001 by unpaired *t*-test (A); ns, non-significant. All error bars are the SEM.

**Supplementary Figure 3. Constitutive activation of Rab21 has no obvious effect on tissue homeostasis, while its loss affects cell survival** (A) Inversed binary images of

*Drosophila* guts stained with DAPI. Cell density was quantified in *N*=3 independent experiments. (B) Immunostaining for the cell junction marker armadillo (arm). *N*=3 independent experiments (C) RT-qPCR of the pro-apoptotic gene *Hid.* *N*=4 independent experiments. $^{**}P < 0.01$ by unpaired *t*-test (A, C); ns, non-significant. All error bars are the SEM.

**Supplementary Figure 4. Rab21 is important for proper endosomal trafficking in enterocytes** (A) PtdIns(3)P was detected in *Drosophila* enterocytes by expressing GFP:2×FYVE. GFP:2×FYVE$^{+}$ dots were quantified in *N*=3 independent experiments. (B) Early endosomes were detected by immunostaining for the early endosomal marker Hrs. Hrs$^{+}$ dots were quantified in *N*=3 independent experiments. (C) PtdIns(3,5)$P_2$ was detected in *Drosophila* enterocytes by expressing mCherry:ML1N2×. Quantification of mCherry:ML1N2×$^{+}$ dots was based on *N*=3 independent experiments. (D) Late endosomes were detected by immunostaining for Rab7. Rab7$^{+}$ dots were quantified in *N*=3 independent experiments. (E) Lysosomes were detected in *Drosophila* enterocytes by expressing GFP:LAMP. Images are representative of *N*=4 independent experiments. $^{*}P < 0.05$; $^{**}P < 0.01$; $^{***}P < 0.001$; $^{****}P < 0.0001$ by unpaired *t*-test (A) or Mann-Whitney U test (A, B, C, D); ns, non-significant. All error bars are the SEM.

**Supplementary Figure 5. Depletion of proteins related to different steps in membrane trafficking do not all phenocopy Rab21 depletion** (A) Quantification of pH3$^{+}$ cell percentages after depletion of the indicated proteins in *N*=3 independent experiments, except for AP-2$^{RNAi}$ (*N*=1). Gene names are highlighted in colors corresponding to the membrane trafficking steps they affect (defined in Figure 5A). (B) The upd3-LacZ staining intensity after depletion of the indicated proteins was quantified in *N*=3 independent experiments, except for AP-2$^{RNAi}$ (*N*=1). Gene names are highlighted as in Figure 5A. *$P$ < 0.05; **$P$ < 0.01; ***$P$ < 0.001 by Mann-Whitney U test (A, B); ns, non-significant. All error bars are the SEM.

**Supplementary Figure 6. Rab21's regulatory effects on autophagy and EGFR signaling are not epistatic in enterocytes** (A) To assess the effects of Rab21 depletion on autophagic flux, we immunostained for ref(2)p. Quantification of ref(2)p$^{+}$ dots was based on *N*=3 independent experiments. (B) TEM of *Drosophila* gut tissue sections. Images are representative of 22 images acquired in *N*=2 independent experiments. The left panels are magnifications of the areas outlined by dotted lines in the right panels. (C) Immunostaining for dpERK after depletion of autophagy-related genes. *N*=2 independent experiments. ****$P$ < 0.0001 by unpaired *t*-test (A); ns, non-significant. All error bars are the SEM.

Supp. Figure1

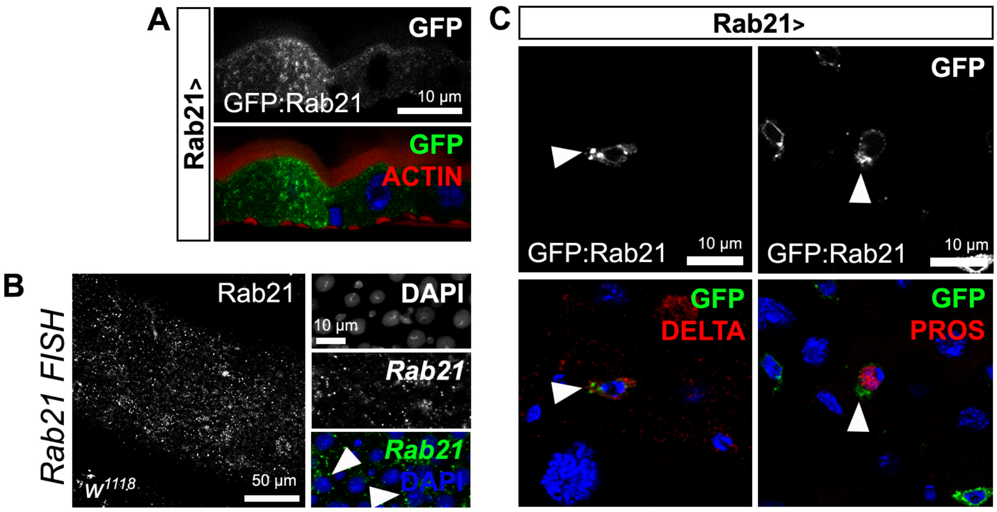

Supp. Figure2

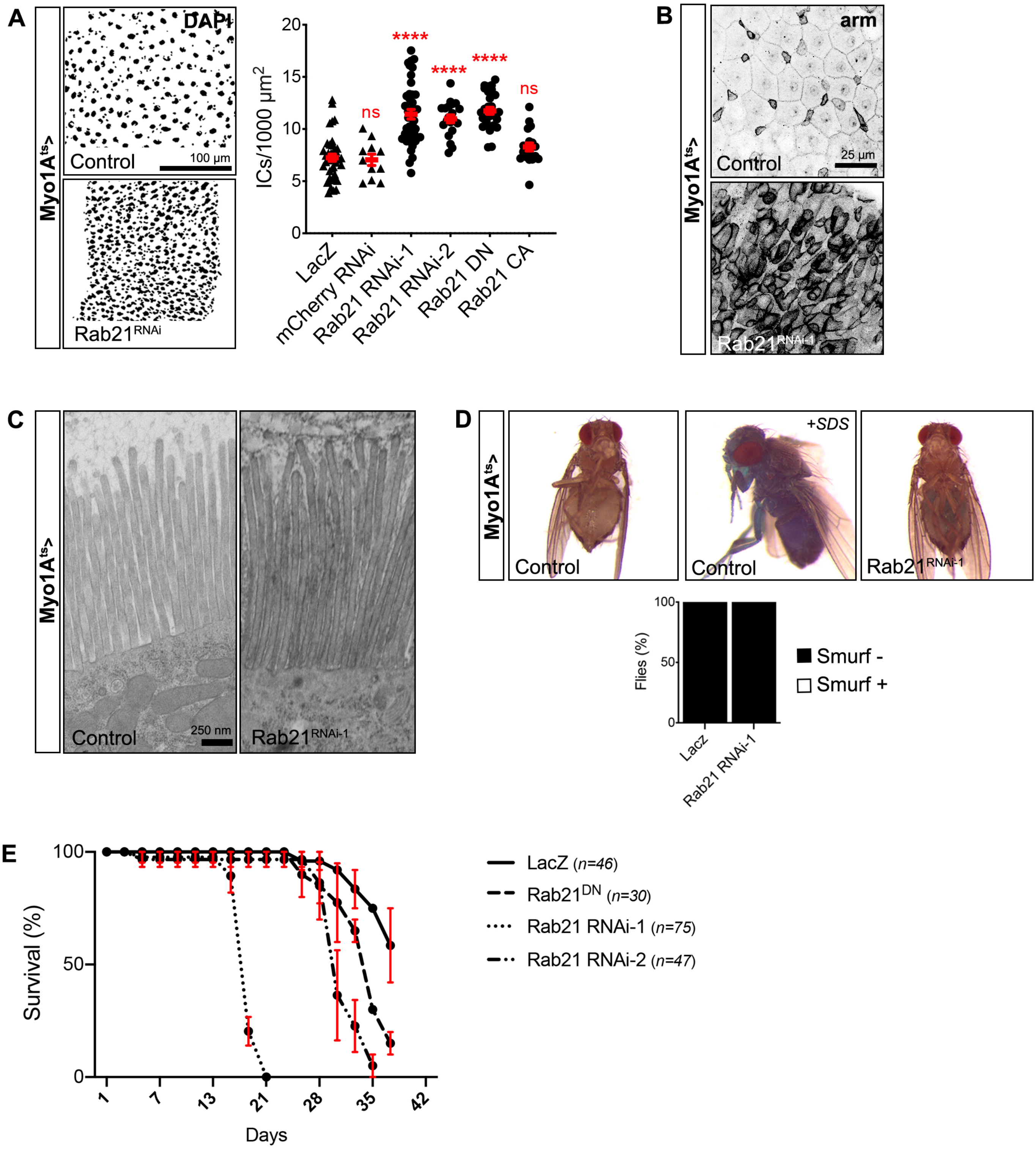

### Supp. Figure3

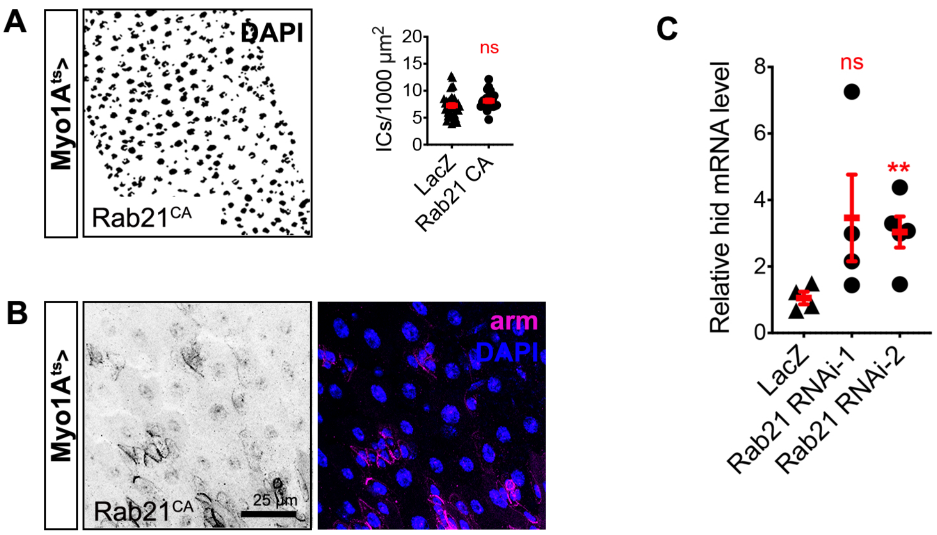

Supp. Figure4

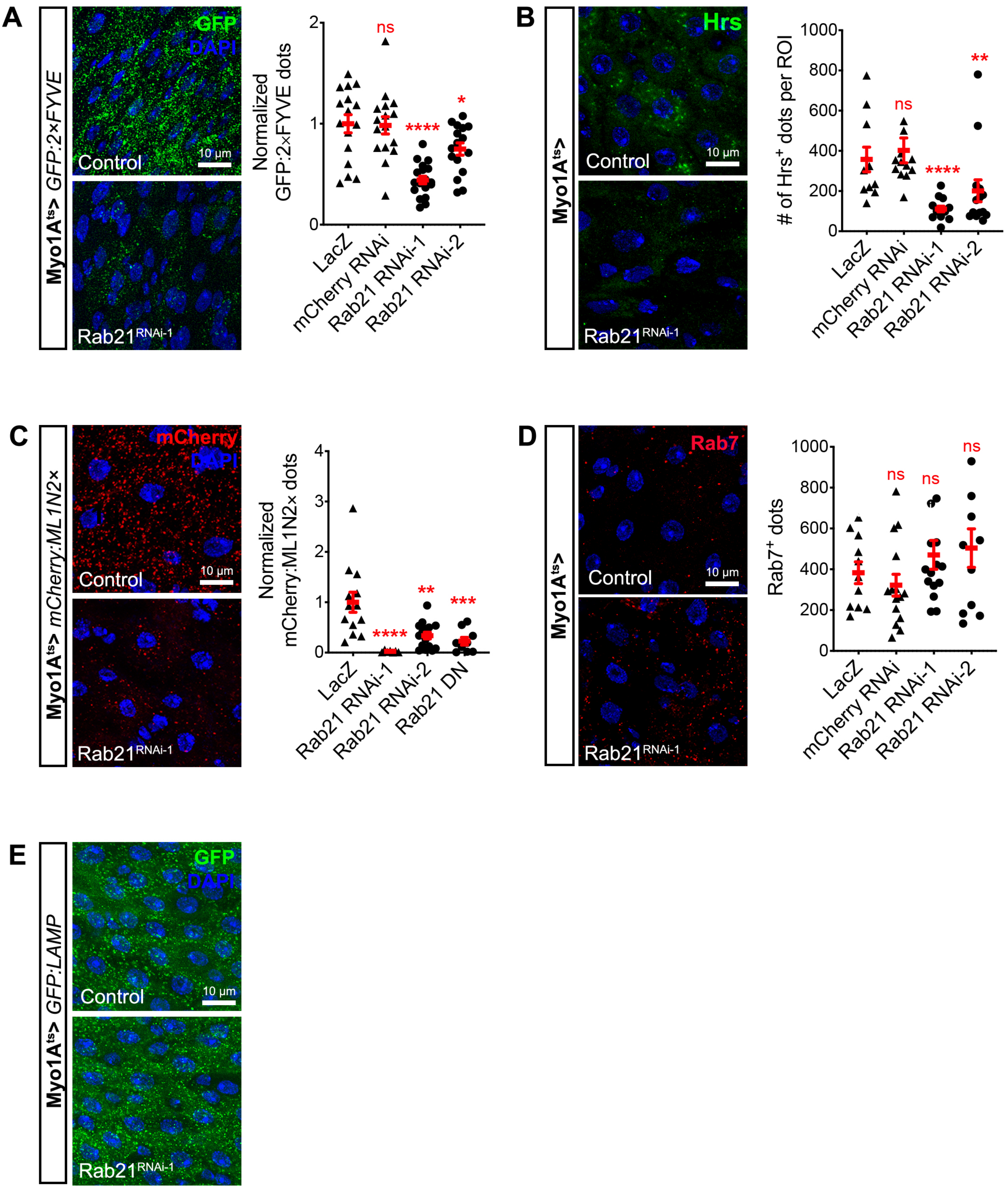

Supp. Figure5

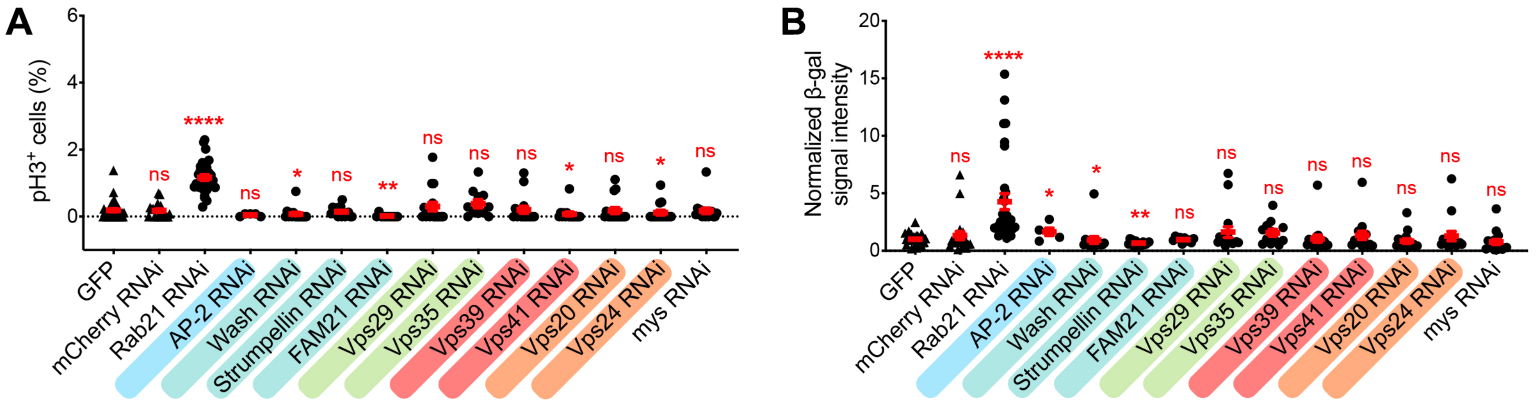

Supp. Figure6

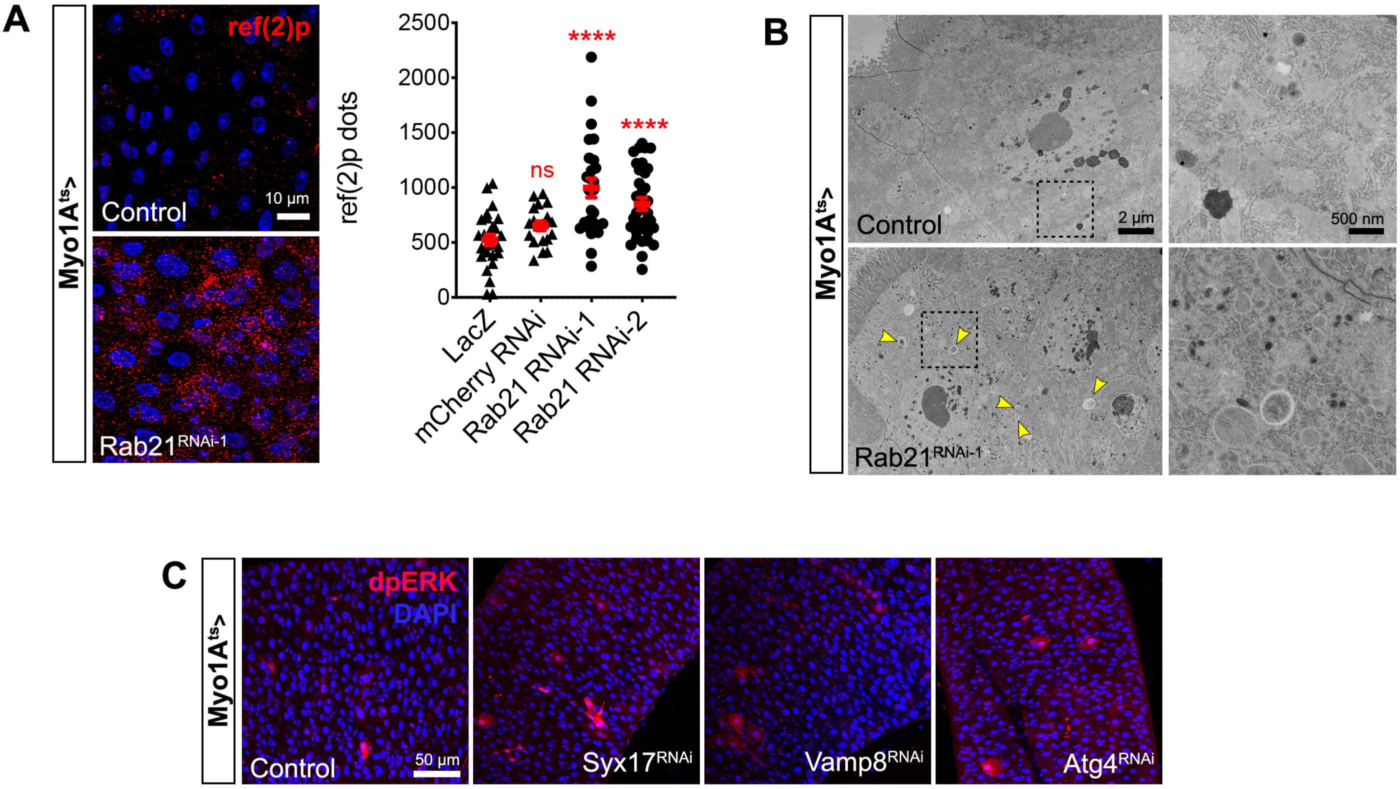
